# Once upon a time in the far south: Influence of local drivers and functional traits on plant invasion in the harsh sub-Antarctic islands

**DOI:** 10.1101/2020.07.19.210880

**Authors:** Manuele Bazzichetto, François Massol, Marta Carboni, Jonathan Lenoir, Lembrechts Jonas J, Rémi Joly, David Renault

## Abstract

**Aim:** Here, we aim to: (i) investigate the local effect of environmental and human-related factors on alien plant invasion in sub-Antarctic islands; (ii) explore the relationship between alien species features and their dependence on anthropogenic propagule pressure; and (iii) unravel key traits conferring invasiveness in the sub-Antarctic.

**Location:** Possession Island, Crozet archipelago (French sub-Antarctic islands).

**Taxon:** Non-native vascular plants (Poaceae, Caryophyllaceae, Juncaceae).

**Methods:** Single-species distribution models were used to explore the effect of high-resolution topoclimatic and human-related variables on the occurrence of six of the most aggressive alien plants colonizing French sub-Antarctic islands. Furthermore, the interaction between alien species traits and their response to anthropogenic propagule pressure was analysed by means of a multi-species distribution model. This allowed identifying the features of species that were associated to low dependence on human-assisted introductions, and were thus potentially more invasive.

**Results:** We observed two main invasion patterns: low-spread species strongly dependent on anthropogenic propagule pressure and high-spread species limited mainly by harsh climatic conditions. Differences in invasiveness across species mostly related to their residence time, life history and plant height, with older introductions, perennial and low-stature species being more invasive.

**Main conclusions:** The availability of high-resolution data improved our understanding of the role of environmental and human-related factors in driving alien species distribution on sub-Antarctic islands. At the same time, the identification of alien species features conferring invasiveness may help anticipating future problematic invasions.

## 1. Introduction

Sub-Antarctic islands, archipelagos scattered within the 54-48°S latitudinal ring, are extremely remote territories which harbour a unique biodiversity with a high degree of endemism (Shaw, 2013). As a consequence of their relatively recent discovery and environmental harshness, these islands have long remained pristine and largely free of human disturbances. Yet, due to the gradual relaxation of these natural barriers, sub-Antarctic islands are now listed among the most threatened environments on Earth. In particular, invasion by alien plants, boosted by ongoing climate changes and increasing human disturbances (Duffy & Lee, 2019; Hughes et al., 2019), has become one of the main threats to the endemic biodiversity of these territories, and is bound to rise in the next decades (Lebouvier et al., 2011; Hughes, Pertierra, Molina-Montenegro, & Convey, 2015). Over the past century, alien plants have been increasingly introduced in the sub-Antarctic region (Frenot et al., 2005; Huiskes et al., 2014). European whalers and scientific activities, respectively in the 19^th^ and 20^th^ century, determined the first main introduction events (Convey & Lebouvier, 2009; Shaw, 2013). Later on and since the mid-twentieth century, climate warming, strong changes in precipitation regimes and the widespread impacts of non-native vertebrates have progressively favoured the establishment of cold-tolerant alien plants on sub-Antarctic islands (Shaw, 2013; Pertierra et al., 2017; Duffy & Lee, 2019). Nevertheless, despite their demonstrated impacts on native biodiversity, little attention has been given to plant invasions compared to animal invasions on these islands (le Roux et al., 2013), leaving a knowledge gap in the mechanisms underpinning plant invasion processes in these unique environments (Greve, Mathakutha, Steyn, & Chown, 2017).

The outcome of any biological invasion is jointly determined by propagule pressure (i.e. frequency of propagules introduction), abiotic conditions (i.e. physico-chemical features of the invaded environment) and biotic features (i.e. alien species characteristics and interactions with the recipient community), with anthropogenic disturbances affecting all three (Richardson & Pyšek, 2006; Catford, Jansson, & Nilsson, 2009; Lembrechts et al., 2016). The relative importance of these factors is, however, context-dependent and species-specific (Catford et al., 2009). In sub-Antarctic islands, due to the high specialization but low diversity of the native flora, biotic interactions are thought to play a minor role (le Roux et al., 2013; Duffy et al., 2017; Moser et al., 2018), so it is mainly the first two factors that determine the distribution and spread of alien plants. First, invasions depend on human-induced propagule pressure: the frequency of propagule introduction correlates with the number of ship landings and is highest in the vicinity of human facilities (Huiskes et al., 2014). Second, local abiotic conditions are strongly limiting, and particularly the climatic mismatch between the conditions prevailing within the alien species’ native range and the conditions prevailing in the sub-Antarctic can strongly constrain invasions (Frenot et al., 2005). Some alien plants are more limited during the introduction phase, while others quickly become relatively independent of human-related propagule-pressure and seem only climatically limited. Once established, the species which are the least dependent on continuous introductions are the most likely to spread widely and become invasive (Richardson & Pyšek, 2006; Catford et al., 2009). Therefore, quantifying the degree of alien species dependence on propagule pressure might aid at identifying potentially invasive species.

A lower dependence on human-related propagule pressure is potentially related to certain species features which are more generally known to affect invasiveness. First of all, alien species with longer residence times are more likely to become independent of anthropogenic propagule pressure (Wilson et al., 2007; Pyšek et al., 2015). Second, certain plant traits are considered key for profiling successful invaders (Pyšek & Richardson, 2008): invasive alien plants across most environments are growing faster and taller than non-invasive alien species, and typically produce resource-acquisitive leaves and many small seeds (van Kleunen, Weber, & Fischer, 2010; van Kleunen, Dawson, & Maurel, 2015). More specifically, Mathakutha et al. (2019) performed a first functional comparison between invasive and non-invasive alien species colonizing the sub-Antarctic Marion Island, reporting that species generally considered invasive had lower plant height, smaller leaf area, lower frost tolerance and higher specific leaf area than other alien species. Nevertheless, it is still unclear which traits can actually make some alien plants less dependent on human-related propagule pressure, and thus more likely to become invasive, especially in the sub-Antarctic islands. This knowledge could facilitate the early screening of highly invasive alien plant species in these environments (Frenot et al., 2005; Mathakutha et al., 2019).

Correlative species distribution models (SDMs) are statistical tools that model the species-environment relationship relying on geo-referenced occurrence data and spatial environmental layers (Guisan, Thuiller, & Zimmermann, 2017). Such models already proved to be valuable tools for analysing alien plant invasion in Antarctica and the sub-Antarctic regions. For instance, Pertierra et al. (2017) modelled the distribution of *Poa annua* and *Poa pratensis* in the Antarctic peninsula as a function of bioclimatic variables, while Duffy et al. (2017) generated future scenarios of invasion across Antarctica and the sub-Antarctic regions using climate-based SDMs. Whilst these previous SDM applications have revealed large-scale determinants of alien plant invasion in the Antarctic biogeographic region, they have up till now failed to account for how environmental and anthropogenic factors regulate alien plant distributions at a spatial resolution that is meaningful for local management. This is chiefly due to the lack of high-resolution environmental (e.g. climatic, topographic) and human-related data layers, which limits the implementation of SDMs at fine spatial resolutions in remote areas (Gutt et al., 2012). A more general limitation inherent to the use of SDMs for modelling biological invasion is that SDMs allow mapping into the geographical space only a snapshot of the current alien species-environment relationship net of dispersal and biotic constraints (i.e. realized distribution), while necessarily underestimating the actual area potentially suitable to a species for establishing and maintaining a viable population (i.e. potential distribution; see Jiménez-Valverde et al., 2011 and Srivastava, Lafond, & Griess, 2019).

The sub-Antarctic Possession Island constitutes an ideal arena to analyse alien plant invasions in the sub-Antarctic region. The availability of historical vegetation observations allows retracing the invasion history of most alien plant species on the island. Moreover, this island witnessed past human colonization and climate changes comparable to the other sub-Antarctic islands, allowing inference on the mechanisms underpinning alien plant invasion in these unique areas. Previous work showed that there is considerable variation in the spread of alien plants established on Possession Island, with some species clustering close to their introduction locations and others spreading widely and far from the initial introduction sites (Frenot et al., 2005), which allows testing for differences in the dependence on human introductions. In the present study, we model the distribution of the most relevant alien plant species colonizing Possession Island using a combination of environmental and human-related spatial data derived at an unprecedented high spatial resolution (i.e. 30-m) for these latitudes that we related to long-term monitoring observations of plant occurrences. Our aim is to test the local effect of environmental and anthropogenic factors on alien plant invasion in sub-Antarctic ecosystems. We hypothesise that both abiotic and human-related factors jointly define the local occurrence of alien plant species, but that these two factors will not be equally important among species. Furthermore, to identify plant characteristics conferring high invasiveness in sub-Antarctic ecosystems, we investigate how plant functional traits affect species dependence on anthropogenic propagule pressures. In this regard, our working hypothesis is that the most invasive species share specific functional characteristics allowing them to become independent of human-assisted introductions and spread widely once established.

## 2. Materials and methods

### 2.1 Study area

The study was carried out on Possession Island in the Crozet archipelago, which is included in the *Réserve naturelle nationale des terres australes françaises* (RNN TAF) and listed as UNESCO World Heritage site since 2019. Possession Island (Figure 1) is characterized by a complex topography, with an altitudinal gradient ranging from 0 to 934 m above the sea level (*Pic du Mascarin*) over a relatively short spatial extent (147 km^2^). The island is characterized by a typical sub-Antarctic climate, with mean annual temperature of 5.6 °C and annual precipitation of 2,300 mm (Météo France, data: 1960-2019). Frequent and strong western winds occur throughout most of the year.

**Figure 1.**
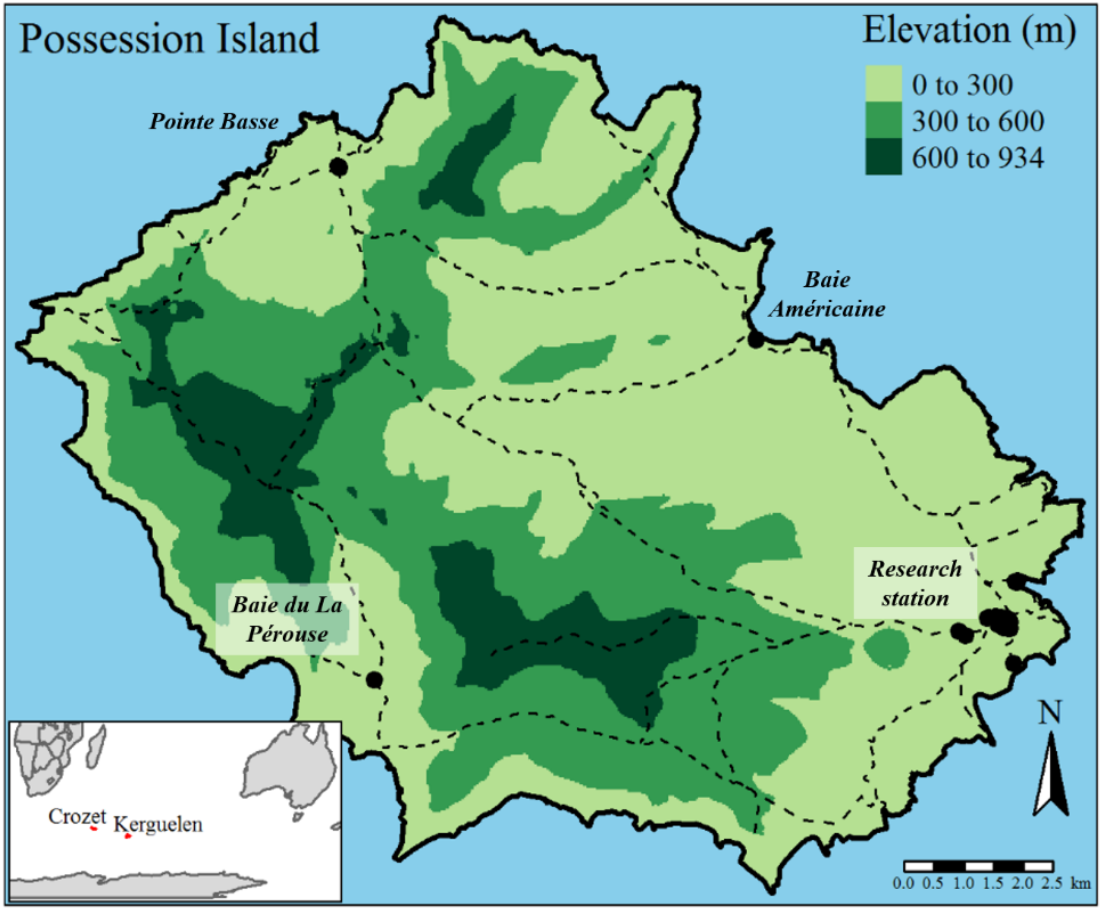
Map of Possession Island showing: gross topography using three altitudinal belts (0-300 m; 300-600 m; 600-934 m); human settlements (black dots); and hiking paths (dashed lines). The inset map reports the geographical location of French sub-Antarctic islands, which include Crozet and Kerguelen archipelagos.

The first human settlements date back to the 19^th^ century, when whalers and sealers established on the north-east side of the island during the hunting season, facilitating a first series of alien species introductions. In 1963, a permanent research station (*Alfred Faure*, hereafter the ‘research station’) was built on the easternmost area of Possession Island, fostering a new invasion front. Beyond the research station, other shelters (inhabited for short periods) are currently present on each side of the island: north (*Pointe Basse*); south-west (*Baie du La Pérouse*); and north-east (*Baie Américaine*). Among these, the research station is by far the biggest human settlement and main hub of propagule introduction. The vegetation at Possession Island has experienced relatively low grazing pressure from sheep in the past (Convey & Lebouvier, 2009), in comparison to other sub-Antarctic islands where introduced large herbivores still strongly affect the distribution of alien plants (Shaw, 2013).

### 2.2 Study species

Despite the 68 alien species recorded on Possession Island (see page 99 of the RNN TAF management plan 2018-2027: https://taaf.fr//content/uploads/sites/2/2019/09/180607-Volet-A_pour-CNPN.pdf), only few have established persistent populations (Frenot, Gloaguen, Massé, & Lebouvier, 2001). In this study, we restricted our analysis to those alien plants that are either known to be generally widespread on sub-Antarctic islands or are particularly widespread on Possession Island, and for which sufficient occurrence data were available (total number of presences > 100). Specifically, we selected the following six species from three different families: *Poa annua* and *Poa pratensis* (Poaceae); *Cerastium fontanum, Sagina procumbens* and *Stellaria alsine* (Caryophyllaceae); and *Juncus bufonius* (Juncaceae). The two grasses, *P. annua* and *P. pratensis*, have colonised most of the sub-Antarctic islands (Shaw, 2013) and are the longest-established alien plants in the Antarctic Peninsula (Pertierra et al., 2017). *Cerastium fontanum* and *S. procumbens* are currently widely distributed in this environment (Frenot et al., 2005; Shaw, 2013) with, in particular, *S. procumbens* exhibiting the highest rate of spread among the alien plants of Marion and Prince Edward Islands (le Roux et al., 2013). Finally, both *J. bufonius* and *S. alsine* currently occur at significant distances from the research station on Possession Island (Frenot el al., 2001). While the former has been recently observed up to the Maritime Antarctica latitudes (Cuba-Díaz, Fuentes, & Rondanelli-Reyes, 2015), the latter has been singled out by some authors as the potentially most problematic future invasive plant species on Possession Island (Frenot et al., 2001; Convey, Key, & Key, 2010).

#### 2.2.1 Species distribution data

We analysed the invasion patterns of the six selected alien plant species relying on georeferenced occurrence (presence/absence) data collected within the context of a yearly vegetation monitoring survey carried out by the RNN TAF since 2010. The vegetation sampling is implemented within a system of 675 squared cells of 100 × 100 m each, where floristic data (presence and abundance of vascular plant species) are collected along with habitat characteristics through phytosociological *relevés* (Dengler, 2016). In this study, we used data collected from 2010 to 2017 (3,354 occurrences for the selected species across 1,572 sampled plots).

#### 2.2.2 Species features and functional trait data

To inform species features (e.g. traits) potentially related to invasiveness, we collected data on plants residence time and functional traits. Residence time positively interacts with propagule pressure in determining plant invasion success (Richardson & Pyšek, 2006; Lockwood, Cassey, & Blackburn, 2005; Pyšek et al., 2015), and this relationship was also observed on sub-Antarctic islands (le Roux et al., 2013; Shaw, 2013; Mathakutha et al., 2019). To test how residence time influences alien species’ dependence on propagule pressure, we considered the introduction date of the selected plants on Possession Island (Frenot et al., 2001) and used this information to assign them to two groups: old *vs* new resident species (Appendix S1, Table S1.1). In particular, we considered as old resident species those which were firstly observed on Possession Island before the research station was built (1963), while referring to the others as new resident species.

We then collated data on seven plant traits commonly used to synthesize species strategies known to be related to invasiveness (van Kleunen et al., 2010; van Kleunen et al., 2015): (1) life history (annual *vs* perennial); (2) plant height; (3) leaf area; (4) specific leaf area (SLA); (5) vegetative reproduction (present *vs* absent, i.e. sexual and vegetative *vs* only sexual reproduction); (6) seed dry mass; and (7) number of seeds per plant. We excluded traits related to flowering since pollinating insects are absent from almost all sub-Antarctic islands (Convey et al., 2010). Life history, plant height and leaf area relate to plant persistence and tolerance to environmental stress (Cornelissen et al., 2003; Pérez-Harguindeguy et al., 2013). In addition, life history is used to assess maximum lifespan and plant height is associated with competitiveness for light and whole plant fecundity (Pérez-Harguindeguy et al., 2013). Specific leaf area is the one-sided leaf area per leaf mass and is associated with resource acquisition and photosynthetic rate (Pérez-Harguindeguy et al., 2013). Reproduction strategy, seed dry mass and number of seeds per plant do not only relate with species persistence, but also with dispersal capacity (Ottaviani et al., 2020). In particular, alien species reproducing predominantly sexually may benefit from lower dispersal limitation and greater genetic diversity (van Kleunen et al., 2015). At the same time, while small and light seeds are better dispersed at longer distances, large-seeded plants may benefit from more stored resources (van Kleunen et al., 2015).

Functional trait data collected in areas environmentally analogous to Possession Island were compiled from the literature (other sub-Antarctic islands, Frenot et al., 2005; Marion Island, Mathakutha et al., 2019). Whenever we could not find information collected in comparable environments, we relied on functional trait data included in the TRY database (Kattge et al., 2020). For each alien species, the dominant reproduction strategy in the study area was assessed relying on expert-based knowledge (personal communication, Lebouvier, M., & Bittebiere, A.K.). Species-specific values of the functional traits are reported in table S1.1 (Appendix S1) along with literature sources.

### 2.3 Topoclimatic layers

To model the species-environment relationship at fine spatial resolution, we first downloaded coarse-grained temperature (BIO1, BIO5 and BIO6 – annual mean temperature, max temperature of the warmest month and min temperature of the coldest month) and annual precipitation (BIO12) grid layers at 1-km resolution (at the equator) from the CHELSA database (Karger et al., 2017) and then disaggregated their spatial resolution using physiographically informed models fitted through geographically weighted regression (GWR; Fotheringham & Rogerson, 2008). This downscaling technique allows statistically predicting the local value of the coarse-grain CHELSA climatic variables as a function of environmental grid layers available at finer spatial resolution (in this study 30-m at the equator, hereafter 30-m) and known to drive microclimate heterogeneity (Lenoir, Hattab, & Pierre, 2017; Lembrechts et al., 2019). GWR-derived topoclimatic layers, beyond allowing to model the species-environment relationship at a more meaningful spatial resolution, have already proved to better account for the complex interactions between macroclimate and topography (Lenoir et al., 2017; Lembrechts et al., 2019).

As using BIO5 (max temperature of the warmest month) and BIO6 (min temperature of the coldest month) in place of BIO1 did not improve species distribution models, we ultimately used BIO1 (hereafter mean temperature) and BIO12 as topoclimatic predictors. A full description of the downscaling procedure is reported in Appendix S2 along with the results of the GWR models.

### 2.4 Human-related layers

As human disturbances are known to favour the establishment of alien plants through propagule introduction and alteration of habitat conditions, we generated a 30-m resolution layer reporting the distance between each human settlement (the research station, *Baie du La Pérouse, Pointe Basse* and *Baie Américaine*) and any location on the island. Specifically, assuming that human disturbance is stronger in more accessible areas, we derived a least cost distance grid layer providing a measure of accessibility. Terrain slope changes between both orthogonally and diagonally neighbouring raster cells were used to compute the cost of reaching any location on Possession Island starting from any human settlement and following all potential paths of raster cells (function “accCost”, “gdistance” R package; Etten, 2018). High costs were thus associated with locations not easily reachable from human settlements due to high topographic roughness (Appendix S3, Figure S3.2).

A network of hiking paths has been designed by RNN TAF to restrict human movements for wildlife conservation purposes, and walking these paths currently constitutes the only authorized way to move across the island. As humans are a critical vector of propagule introduction and dispersal on sub-Antarctic islands, we derived a 30-m resolution raster layer reporting the distance between any location on Possession Island and the closest hiking path using the function “distance” from the “raster” R package (Hijmans, 2019) (Appendix S3, Figure S3.2).

### 2.5 Alien species distribution modelling

The occurrence probability of the six studied alien plant species was separately modelled as a function of the topoclimatic (mean temperature and annual precipitation) and human-related variables (path distance and least cost) by means of logit binary generalized linear models (GLM). The single-species distribution models (single-SDMs) were trained and tested on datasets obtained through a re-sampling procedure of the presence/absence data performed in the environmental space to reflect all available environmental conditions on Possession Island (Lenoir et al., 2010; Hattab et al., 2017; see Appendix S4). All four topoclimatic and human-related predictors were retained to fit the single-SDMs as the relative variance inflation factor (function “vif”, R package “car”; Fox & Weisberg, 2019) was always below a threshold of 3. Second-order polynomial terms were included in the model to allow for intermediate niche optima of the species or in case lack-of-fit tests detected consistent departure from linearity in the profile of Pearson residuals (function “residualPlots”, R package “car”; Fox & Weisberg, 2019). The statistical significance of each predictor was tested using type II analysis of deviance (function “Anova”, R package “car”; Fox & Weisberg, 2019). We then computed the likelihood profile-based 95% confidence intervals of the regression parameters.

Single-SDMs predictive performance was measured using the true skill statistic (TSS, equal to sensitivity + specificity – 1; function “ecospat.max.tss”, R package “ecospat”; Broennimann, Di Cola, & Guisan, 2018) computed on the testing datasets obtained through the environmental matching described in Appendix S4. We used the TSS as it has desirable properties of other accuracy measures (e.g. Kappa and AUC), while being unaffected by prevalence (Allouche, Tsoar, & Kadmon, 2006). Also, we computed the deviance-based R^2^ value as a measure of goodness-of-fit of each single-SDM.

The occurrence probability estimated by the full single-SDMs (including both topoclimatic and human-related predictors) for each alien plant species was mapped on a 30-m raster grid layer to visualise their predicted distribution across Possession Island.

### 2.6 Relationship between plant traits and alien species dependence on propagule pressure

As preliminary analyses, we measured the relative importance of human-related variables in determining alien species occurrence in the single-SDMs. To this aim, we used the sum of Akaike weights (*w*), which provides an easily interpretable measure of variable importance (it ranges from 0 to 1, with a high value for a given variable indicating its high importance relative to the others; Burnham & Anderson, 2002). Then we graphically related the species-specific values of the functional traits to the sum of weights to look for relationships between plant traits and the importance of human-related variables (see Appendix S7).

Secondly, we investigated how the interaction between human-related variables and plant traits affected alien species occurrence in a multi-species distribution model (multi-SDM), focusing on those functional traits that showed some relationship with the dependence on human-related variables in the single-SDMs. To this aim, we modelled the occurrence of all alien species together as a function of topoclimatic and human-related variables by means of a logit binary GLM, including the interaction between species identity and topoclimatic variables on the one hand and the interaction between species functional traits and human-related variables on the other hand. This allowed exploring how the effect of human-related variables on alien species occurrence varied according to plant traits, while controlling for species-specific responses to topoclimate. To select the most parsimonious model, we fitted all possible sub-models including different combinations of the functional traits-anthropogenic variables interaction terms (function “dredge”, R package “MuMIn”; Barton, 2019), always retaining the species-topoclimate interaction terms and the main effect of path distance and least cost in each candidate sub-model. Then, we computed the sum of Akaike weights for each model term and used the evidence ratio as a measure of the relative importance of variables (Massol et al., 2007; Burnham & Anderson, 2002). Specifically, we computed the evidence ratio of the *i-th* variable (*ER*_*i*_) as the odds of its sum of Akaike weights:

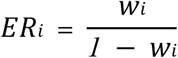

The evidence ratio was then compared with its expected value (*ER*_*null*_) under the “null hypothesis” that the variable explained as much deviance as a randomly generated explanatory variable, and would thus be as likely as not to be incorporated in the best models. As all the variables were tested in a balanced design, *ER*_*null*_ = 1 in all tested cases. Following Massol et al. (2007), the effect of a variable *i* was deemed unlikely if *ER*_*i*_ < 0.37 × *ER*_*null*_, implausible when 0.37 × *ER*_*null*_ < *ER*_*i*_ < *ER*_*null*_, plausible when *ER*_*null*_ < *ER*_*i*_ < 2.72 × *ER*_*null*_, and likely when *ER*_*i*_ > 2.72 × *ER*_*null*_. These thresholds correspond to differences in Akaike information criterion equal to +2 or −2, which are commonly admitted as a good rule-of-thumb gap to compare model performance.

## 3. Results

### 3.1 Effect of topoclimatic and human-related variables on single species distribution

Predictive performances of the single-SDMs varied greatly across species (Table 1): high values of TSS were observed for *P. pratensis, S. alsine* and *J. bufonius* (0.80-0.82), while low values were obtained for the remaining species (from 0.09 to 0.29). The R^2^ values showed a similar trend, with the highest value obtained for *J. bufonius* (0.48) and the lowest for *C. fontanum* (0.02).

**Table 1.** Single-SDM type II analysis of deviance tables and performance measures (R^2^ and TSS). LR: Likelihood Ratio statistic; Df: degrees of freedom; p-val: p-value (*** p < 0.001; ** p < 0.01; * p < 0.05).

|  | *Poa pratensis* |  |  | *Juncus bufonius* |  |  | *Stellaria alsine* |  |  | *Poa annua* |  |  | *Sagina procumbens* |  |  | *Cerastium fontanum* |  |  |
| --- | --- | --- | --- | --- | --- | --- | --- | --- | --- | --- | --- | --- | --- | --- | --- | --- | --- | --- |
| Predictors | LR | Df | p-val | LR | Df | p-val | LR | Df | p-val | LR | Df | p-val | LR | Df | p-val | LR | Df | p-val |
| Mean temperature | 5.083 | 1 | * | 32.516 | 2 | *** | 0.312 | 1 | = 0.576 | 55.538 | 1 | *** | 62.554 | 2 | *** | 18.325 | 2 | *** |
| Annual precipitation | 106.443 | 1 | *** | 73.647 | 1 | *** | 68.406 | 2 | *** | 4.759 | 1 | * | 3.681 | 1 | = 0.055 | 0.041 | 1 | = 0.840 |
| Least cost | 10.837 | 1 | *** | 40.250 | 1 | *** | 4.420 | 1 | * | 5.778 | 1 | * | 1.118 | 1 | = 0.290 | 12.483 | 1 | *** |
| Path distance | 15.623 | 1 | *** | 8.740 | 1 | ** | 7.524 | 1 | ** | 3.927 | 1 | * | 6.782 | 1 | ** | 2.877 | 1 | = 0.089 |
| $R^2$ | | | 0.46 | | | 0.48 | | | 0.36 | | | 0.06 | | | 0.10 | | | 0.02 |
| TSS |  |  | 0.80 |  |  | 0.81 |  |  | 0.82 |  |  | 0.19 |  |  | 0.29 |  |  | 0.09 |

Overall, the occurrence of *P. pratensis, S. alsine* and *J. bufonius* appeared to be strongly conditioned by both topoclimatic and human-related variables, while *C. fontanum, P. annua* and *S. procumbens* were less affected by human-related variables (Table 1, Figures 2 and 3).

**Figure 2.**
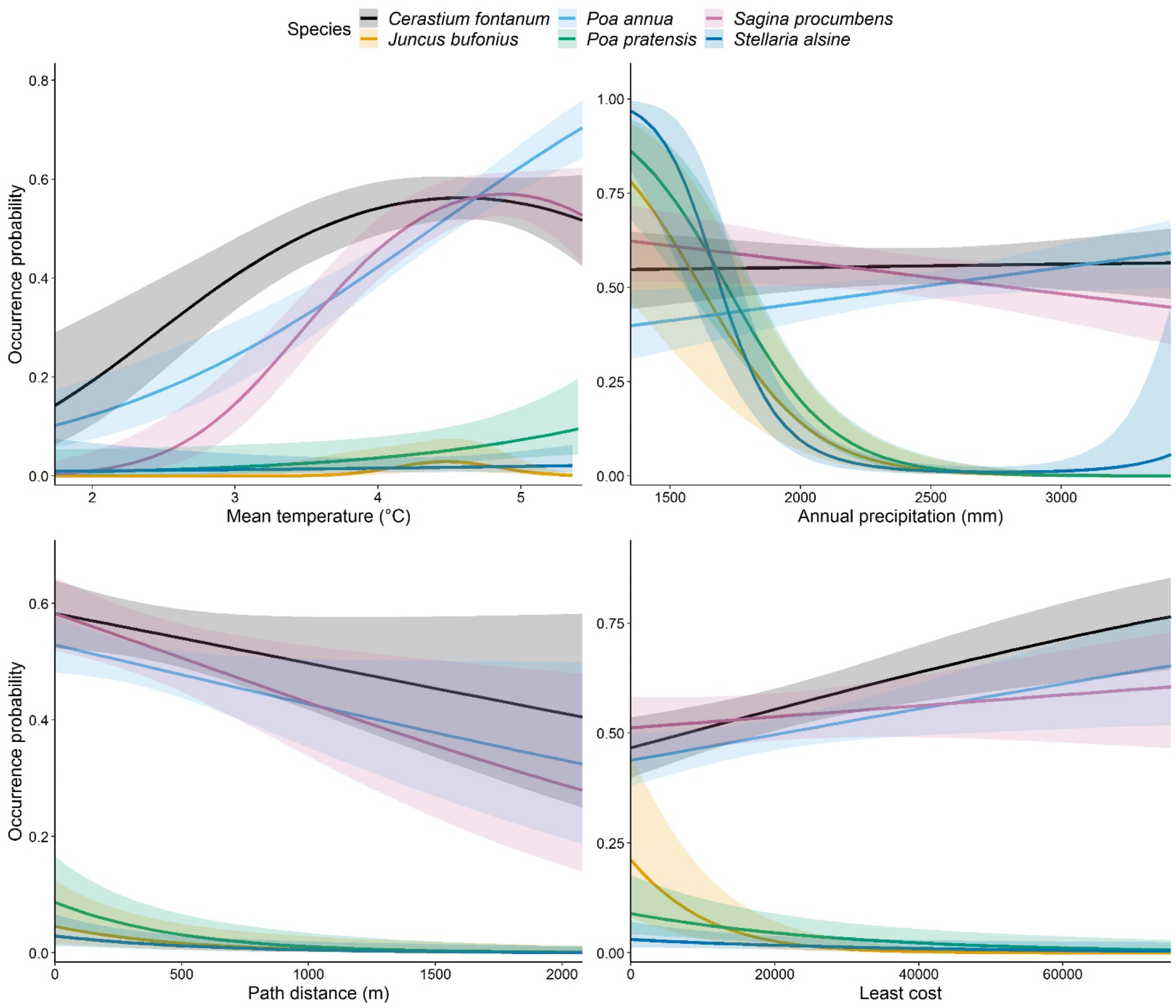
Response curves of the analysed alien species in the single-SDMs.

**Figure 3.**
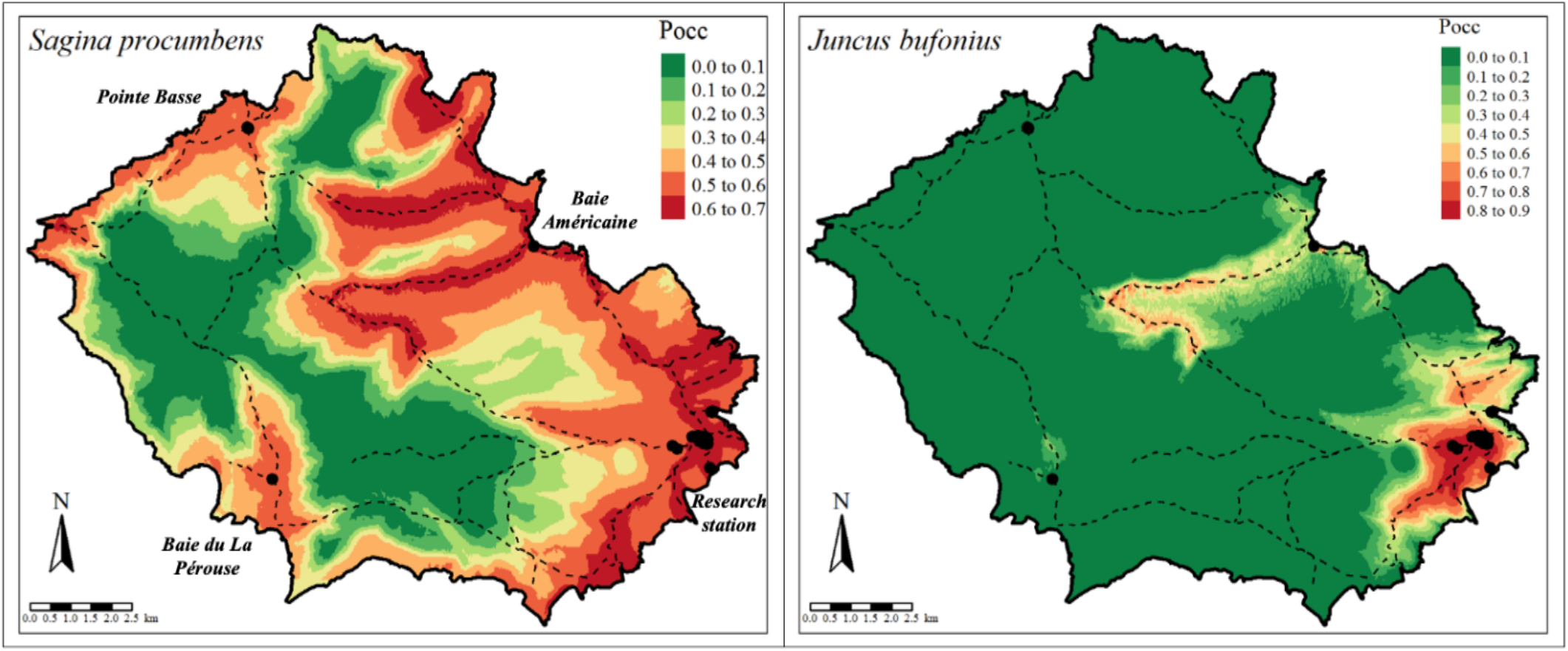
Predicted occurrence of Sagina procumbens and Juncus bufonius. Pocc: occurrence probability. Dashed lines represent hiking paths, while black dots represent human settlements. Occurrence maps of the other alien species are reported in Appendix S6 (Figure S6.4 and S6.5).

All alien species, except *S. alsine*, exhibited a significant positive or humped-shaped relationship with mean temperature (Table 1), meaning that their occurrence probability increased with increasing temperature (Figure 2, Appendix S5). More specifically, the occurrence probability of *J. bufonius, S. procumbens* and *C. fontanum* peaked at mean temperature values around 4.5 °C, while the presence of *P. pratensis* and *P. annua* increased more or less linearly with temperature.

Annual precipitation significantly affected the presence of *P. pratensis, S. alsine* and *J. bufonius*, while it had a minor influence on the occurrence of the other species (Table 1). In particular, the odds of finding *P. pratensis* and *J. bufonius* decreased approximately by 90% for each 500 mm increment in annual precipitation, while the occurrence probability of *S. alsine* sharply decreased for annual precipitation values above 1,500 mm (Figure 2).

All species except *C. fontanum* exhibited a significant negative relationship with path distance (i.e. the occurrence probability of the species decreased at increasing distances from the hiking paths), though its influence varied among species (Table 1, Figure 2, Appendix S5). In this regard, the odds of finding *P. pratensis, S. alsine* and *J. bufonius* decreased respectively by 20%, 16% and 19% moving 100 m away from the paths, while the odds of finding *P. annua* and *S. procumbens* decreased by 4% and 6%, respectively.

Least cost distance to settlements appeared to influence the occurrence of all analysed species except *S. procumbens* (Table 1). In particular, the odds of finding *P. pratensis, S. alsine* and *J. bufonius* decreased, respectively, by 17%, 13% and 44% for each increment of 5,000 units of cost of travelling a given path from a human settlement (Figure 2, Appendix S5). On the contrary, *C. fontanum* and *P. annua* showed a positive relationship with least cost, with their odds of occurring increasing respectively by 9% and 5% for each increment of 5,000 units of cost of travelling a given path from a human settlement (Figure 2 and Appendix S5).

### 3.2 Plant traits and species dependence on propagule pressure

In the preliminary analyses, residence time, life history, vegetative reproduction and plant height showed some relationship with the sum of weights of the human-related variables in the single-SDMs (Appendix S7, Figure S7.6 and S7.8), while seed- and leaf-related traits clearly showed no relationship (Appendix S7, Figure S7.7 and S7.9).

Then, the multi-SDM confirmed significant interactions of residence time, life history and plant height with the human-related variables (Appendix S7, Figure S7.10). Residence time and plant height appeared to interact with both human-related variables, while life history seemed to interact only with least cost in determining alien species occurrence. In particular, the effect of human-related variables on alien species occurrence varied with plant height. For instance, the occurrence probability of taller plants sharply decreased when moving away from human facilities, while a weaker and sometimes opposite trend was observed for plant species of shorter statures (Figure 4a,b and Appendix S7, Figure S7.11). In addition, old residents were on average less affected by the human-related variables than new residents (Figure 4a,b and Appendix S7, Figure S7.11a). Finally, perennials appeared to be on average less negatively affected by least cost distance to human settlements than annuals (Figure 4c and Appendix S7, Figure S7.11b,c).

**Figure 4.**
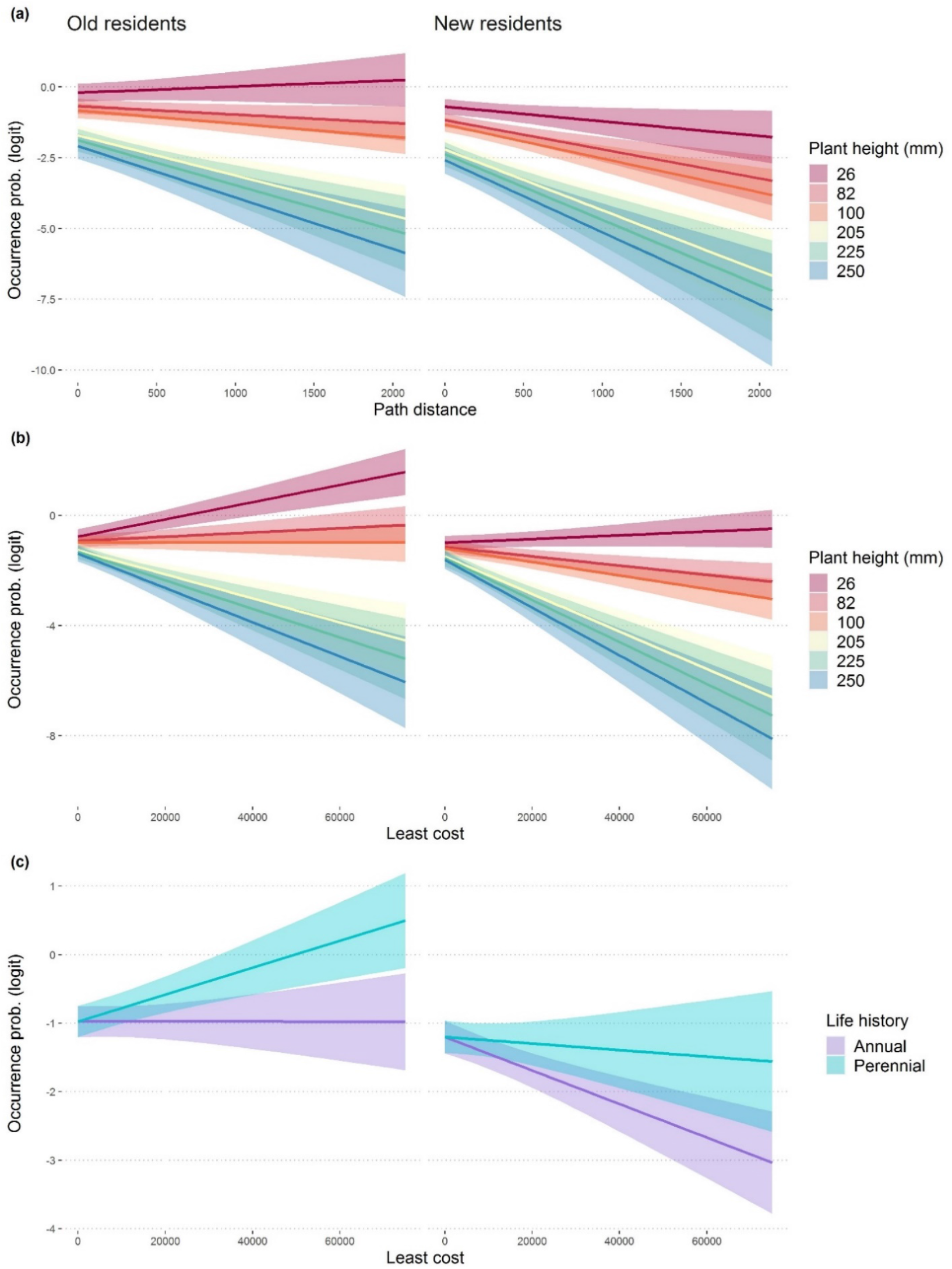
Effect of the interaction between human-related variables and plant features on alien species occurrence probability (logit scale). Panel (a): effect of the path distance-plant height interaction on old and new resident occurrence probability. Panel (b): effect of the least cost-plant height interaction on old and new (annual) resident occurrence probability. Panel (c): effect of least cost distance to human settlements on annual and perennial (100 mm height) alien species occurrence (for both old and new residents). All plots are reported on a logit scale.

## 4. Discussion

As hypothesised, both environmental and human-related variables locally affected alien plant species occurrence on Possession Island, though with differences among the studied species. Overall, results confirmed the key role of human-related propagule pressure in favouring alien plant species establishment and spread on sub-Antarctic islands (Frenot et al., 2005; le Roux et al., 2013; Shaw, 2013), though we also observed a significant effect of abiotic conditions. Indeed, climate barriers seemed to prevent alien plant species occurrence in the most environmentally stressful areas of the island, as found in sub(ant)arctic mountain regions by Lembrechts et al. (2016). In particular, our results suggested the existence of two main invasion patterns arising from the species-specific dependence on human-related propagule pressure (Frenot et al., 2005; Shaw, 2013): low-spread species (*P. pratensis, S. alsine* and *J. bufonius*) strongly relying on human-assisted dispersal along hiking trails and in the vicinity of human settlements; and high-spread species (*C. fontanum, P. annua* and *S. procumbens*) mostly limited by harsh climatic conditions at high altitudes. Differences in plant invasiveness appeared to be influenced by residence time, life history and plant height, with old residents and perennial short species being more invasive.

Due to their dependence on human-related variables, low-spread species were predicted to occur mainly close to hiking paths and human settlements, pointing to the importance of anthropogenic activities as key drivers of continuous propagule pressure favouring species establishment (Whinam et al., 2005; Pickering & Mount, 2010). Once introduced through ship-to-shore transport, propagules are then likely to be dispersed on hiking paths through trampling (Whinam et al., 2005). However, the harsher environmental conditions characterizing the west side of Possession Island also limit the occurrence of low-spread species to the east. In particular, the west-east gradient of annual precipitation (Appendix S2, Figure S2.1) clearly overlaps with the low-spread species distribution (Figure 3 and Appendix S6), suggesting that their establishment might be prevented in areas with abundant precipitation. Nevertheless, the precipitation gradient is also connected to human presence, so that in the western side of the island (less inhabited and more preserved) anthropogenic propagule pressure is weaker. In any case, our results evidenced that low-spread species may lack important adaptations to successfully colonise less disturbed areas with limiting abiotic conditions, while remaining relegated to areas of high human presence.

On the contrary, high-spread species appeared weakly (yet positively) influenced by human-related variables, suggesting that, in spite of the undisputed importance of anthropogenic activities in promoting alien plants establishment (Whinam et al., 2005; Huiskes et al., 2014), high-spread species may possess key traits releasing them from direct dependence on anthropogenic propagule pressure. Consequently, these species appeared to be mostly limited by the extreme climatic conditions of the high and cold inner sectors of Possession Island. This result is in line with the findings from Chwedorzewska et al. (2015), who documented the rapid expansion of *Poa annua* from the Arctowski research base (King George Island) towards wilder areas of the maritime Antarctic Peninsula. Nevertheless, the low predictive performance of high-spread SDMs indicates that the occurrence of these species is poorly explained by the influence of topoclimatic and human-related variables, so that other factors not considered here (e.g. soil properties, plant-soil microbiota interactions, snow cover) may play an important role in driving their distribution at even finer spatial resolutions. In this regard, better performances of the SDMs for high-spread species could have probably been achieved using alien species abundances, which are more informative of the relative habitat suitability than presence-absence data (Howard et al., 2014).

Critically, although we managed to obtain relatively high-resolution topoclimatic data, it is important to realize that the CHELSA climate for the island (1) might lack the accuracy it has at temperate latitudes, being based on extrapolations from a single weather station, and (2) represents air temperature only, while short plants as those analysed here relate more strongly to soil and near-surface temperatures (Convey, Coulson, Worland, & Sjöblom, 2018; Lembrechts et al., 2019). This highlights the need for *in-situ* soil- and near-surface temperature measurements in remote locations to get more ecologically meaningful climate data (Lembrechts et al., 2020).

Although the small set of analysed alien plant species calls for caution in interpretation, we confirmed here that certain plant traits confer greater invasiveness in sub-Antarctic environments. By relating plant traits to species responses to human-related variables and analysing the effect of their interaction on alien species occurrence, we found evidence that low-stature was a key feature that discriminated invasive from non-invasive alien plant species under the harsh abiotic conditions of Possession Island. Noteworthy, residence time and life history also appeared to affect species invasiveness.

As similarly reported by Mathakutha et al. (2019), we found that high-spread species were of shorter statures than low-spread species. Consistently, we observed a sharper decrease in the occurrence probability of taller plants moving away from both hiking paths and human settlements. As plant height is generally associated with species adaptations to harsh environments (Cornelissen et al., 2003) and, specifically, low-stature has been attributed to frost avoidance mechanisms in high mountains (Márquez, Rada, & Fariñas, 2006; Ladinig, Hacker, Neuner, & Wagner, 2013), species of shorter statures may be reasonably favoured in windy and cold sub-Antarctic environments (Mathakutha et al., 2019) and therefore be more easily released of human dependence. Indeed, the importance of functional traits providing tolerance to abiotic stress increases with environmental harshness, even under strong anthropogenic disturbance (Zefferman et al., 2015). Further, our results supported the hypothesis that residence time positively affects invasiveness (Lockwood et al., 2005; Pyšek et al., 2015), though with some exceptions. Generally, old residents (e.g. *C. fontanum* and *P. annua*) were less dependent on human-related propagule pressure and more widely spread than new resident species. Nevertheless, among the old residents, *P. pratensis* was strongly dependent on human-related variables and was still mostly restricted to the original introduction sites. On the other hand, *S. procumbens*, a new resident, has been able to spread extensively and quicker than the other (old residents) high-spread species. However, this might be due to the difference in plant height of the two species: while *P. pratensis* is among the tallest analysed species, *S. procumbens* is the shortest. The multi-SDM showed that perennials were slightly less dependent on human presence than annuals (Figure 4c). Although annuals might benefit from high dispersal abilities (e.g. abundant light seeds) and usually spread quicker and wider (Pertierra et al., 2017), perennials can sustain short growing seasons (Frenot et al., 2001; Shaw, 2013) and potentially colonise harsher environments (Dietz & Edwards, 2006). In our case, short perennials (e.g. *C. fontanum*) might be favoured over tall annuals (e.g. *J. bufonius*) due to the interaction between stress-tolerant traits, such as plant height, and high abiotic tolerance.

Albeit the interaction between vegetative reproduction and human-related variables was not included in the most parsimonious multi-SDM, alien species may still benefit from sexual reproduction, as suggested from the lower importance of human-related variables for the occurrence of alien plants reproducing sexually in the single-SDMs (Appendix S7). As discussed above, the high dispersal potential of sexually reproducing alien species, together with the ability to form rich seed banks (Wódkiewicz et al., 2014), may foster their extensive spread as, for instance, observed for *P. annua* in the Antarctic Peninsula (Pertierra et al., 2017). Nonetheless, by reproducing vegetatively, perennials might outcompete annuals in maintaining viable and persistent populations during unfavourable seasons. Finally, in spite of their acknowledged importance in conferring invasiveness in sub-Antarctic islands (Mathakutha et al., 2019), we found no evidence of the role of seed and leaf traits in affecting species dependence on human-related variables. This is possibly due to the small set of analysed alien species or to the lack of abundance data, which might have prevented the emergence of further functional traits-anthropogenic variables relationships.

Despite some limitations inherent to our dataset (e.g. limited number of species, lack of alien species abundance data) and other typical limitations specific to invasive species distribution modelling (e.g. underestimation of invasion potential, see Jiménez-Valverde et al., 2011), our approach allowed identifying fine-scale drivers of alien plant species distribution, along with the most likely features that favour their spread beyond sources of continuous human-assisted introductions. Combining information on both plant invasiveness and sub-Antarctic islands invasibility, our study provides relevant insights for anticipating future problematic invasions in these remote and unique environments.

## Supporting information

Environmental matching

## Acknowledgements

We would like to thank Marc Lebouvier (Ing. CNRS) for his suggestions which strongly improved the manuscript. We acknowledge the contribution of Lise Chambrin (Ing. RNN TAF) and all civil volunteers of the Institut Polaire Français Paul-Emile Victor (IPEV) who collected data used in this research. Also, we thank Dr. Matos, Prof. Convey and Prof. Hortal for their helpful suggestions and comments, which allowed strongly improving the first version of the manuscript.

## Funding

Project Granted by the French Polar Institute Paul-Emile Victor ‘IPEV 136 Subanteco’, Institut Universitaire de France (IUF) ‘ENVIE’, Stratégie d’attractivité durable (SAD) Région Bretagne ‘INFLICT’. JJL is funded by the Research Foundation Flanders (FWO, project OZ7828), MC by a Rita Levi-Montalcini grant.

Version 3 of this preprint has been peer-reviewed and recommended by Peer Community In Ecology (https://doi.org/10.24072/pci.ecology.100065).

## Authors’ contribution

DR, MB, FM and JL conceived the idea; MB analysed the data with FM, MC and JL; MB led the writing of the manuscript. All authors contributed critically to the drafts and gave final approval for publication.

## Data availability statement

Data and R code available on Zenodo: https://doi.org/10.5281/zenodo.4287498

