## Supplementary material for "Once upon a time in the far south: Influence of local drivers and functional traits on plant invasion in the harsh sub-Antarctic islands"

### Appendix S1

*Table S1.1 – Species-specific values of the functional traits used in the analyses. First obs. date: first observation date of the selected alien plant species along with the corresponding resident group, relative to the building of the research station at Possession Island in 1963; Veg. repr.: vegetative reproduction (present vs absent); SLA: specific leaf area; Seed D.M.: seed dry mass; Seed n°/plant: number of seeds per plant. Superscript letters refer to literature sources: [a] Frenot et al. (2001); [b] Frenot et al. (2005); [c] Lebouvier M. & Bittebiere A.K., personal communication; [d] Mathakutha et al. (2019); [e] Kattge et al. (2020). In particular, superscripts letters posted on the column header indicate that the trait values where all retrieved from a single source, while superscripts letters posted upon specific trait values indicate that a source different from the one appearing on the column header was consulted.*

| *Species* | *First obs. date*[a] | *Resident group*[a] | *Life history*[b] | *Veg. repr.*[c] | *Plant height*[d] *(mm)* | *Leaf area*[d] *(mm²)* | *SLA*[d] *(mm² mg⁻¹)* | *Seed D.M.*[e] *(mg)* | *Seed n°/plant*[e] |
|---|---|---|---|---|---|---|---|---|---|
| *Cerastium fontanum* | 1901 | OLD | Perennial | Abs. | 100.78 | 163.54 | 19.75 | 0.14 | 1290 |
| *Poa annua* | 1906 | OLD | Annual | Abs. | 82.30 | 545.21 | 29.30 | 0.25 | 3665.5 |
| *Poa pratensis* | 1906 | OLD | Perennial | Pres. | 205.45 | 1135.64 | 17.67 | 0.25 | 208 |
| *Sagina procumbens* | 1978 | NEW | Annual | Abs. | 25.71 | 24.27 | 25.07 | 0.02 | 875.5 |
| *Juncus bufonius* | 1984 | NEW | Annual | Abs. | 250[e] | 80.50[e] | 15.10[e] | 0.02 | 34000 |
| *Stellaria alsine* | 1996 | NEW | Perennial | Pres. | 225[e] | 63[e] | 35.20[e] | 0.09 | 201 |

**Appendix S2**

*Downscaling of temperature and annual precipitation CHELSA grid layers*

To obtain high spatial resolution topoclimatic variables, we downloaded coarse-grained (1-km resolution at the equator) temperature (BIO1, annual mean temperature; BIO5, max temperature warmest month; BIO6, min temperature coldest month) and annual precipitation (BIO12) grid layers from the CHELSA database (Karger et al., 2017). Then, we disaggregated their spatial resolution using physiographically informed models fitted through geographically weighted regression (GWR; Fotheringham & Rogerson 2008). Specifically, following the approach described in Lenoir, Hattab, & Pierre (2017) and Lembrechts et al. (2019), the coarse-grained CHELSA grid layers were statistically downscaled as a function of topographic and environmental grid layers available at a finer spatial resolution (in this study 30-m at the equator, hereafter 30-m).

As a first step, we derived a series of topographical and environmental grid layers at 30-m spatial resolution to use as predictors in GWR for downscaling the original CHELSA bioclimatic layers (Table S2.2). A 30-m digital surface model (DSM) across Possession Island was obtained from the Advanced Land Observing Satellite ©JAXA (ALOS; Tadono et al., 2014). The northness (i.e. due south to due north surface direction), eastness (i.e. due west to due east surface direction) and slope were derived from the ALOS DSM using the "terrain" function from the "raster" R package (Hijmans, 2019). Successively, using the "r.sun" tool (Hofierka & Šúri, 2002; GRASS Development Team, 2018), we derived a 30-m insolation time layer reporting the time a given location (here raster cell) is hit by the sunlight on an annual basis. Specifically, the layer was computed averaging the monthly insolation time values (atmospheric turbidity, e.g. optical thickness of the atmosphere, monthly corrected by the corresponding Linke Turbidity Coefficient - data gathered from the SoDa website: http://www.soda-pro.com/). A 30-m NDVI layer was then computed from a Landsat 7 ETM+ scene (scene ID: LE71510922010024SGS00) following Young et al. (2017). Due to the omnipresence of clouds covering Possession Island, we were able to find only one scene (acquired in January 2010) with less than 20% cloud cover, thus usable to accurately compute the NDVI. Lastly, we derived two 30-m grid layers reporting the distance from any location on Possession Island to, respectively, the main waterbodies and the coastline using the function "distance" from the "raster" R package (Hijmans, 2019). The vector maps of the main waterbodies and the shoreline of Possession Island were gathered from OpenStreetMap (OpenStreetMap contributors, 2017).

*Table S2.2 - Descriptive list of the environmental layers used to downscale CHELSA temperature and precipitation layers on Possession Island along with their: sources, range values (min. and max.) and unit of measurement (in brackets). OSM: OpenStreetMap (OpenStreetMap contributors 2017).*

| *Environmental layer* | *Source* | *Range* |
|---|---|---|
| DSM | ALOS | 0 – 909 (m) |
| Slope | DSM-derived | 0 – 82 (deg.) |
| Northness | DSM-derived | -1 – +1 (adim.) |
| Eastness | DSM-derived | -1 – +1 (adim.) |
| NDVI | Landsat 7 ETM+ | -0.5 – +0.54 (adim.) |
| Insolation time | DSM-derived | 0 – 12 (hour) |
| Distance from waterbodies | OSM | 0 – 3.9 (km) |
| Distance from coastline | OSM | 0 – 5 (km) |

As a second step, we fitted a GWR model separately for BIO1, BIO5, BIO6 and BIO12 as a function of the above-mentioned 30-m topographical and environmental layers (function “gwr” from the R package “spgwr”; Bivand & Yu, 2017). In particular, BIO1, BIO5 and BIO6 were regressed on elevation, slope, northness, eastness, NDVI, distance from waterbodies and from the shoreline, while, following Xu et al. (2015), BIO12 was modelled as a function of elevation, slope, northness, eastness and NDVI. The GWR models were fitted using a fixed Gaussian kernel, whose bandwidth was iteratively selected minimizing a cross-validation score (function “bw” from the R package “spgwr”; Bivand & Yu, 2017). The predictive accuracy of the GWR models was then assessed by computing and mapping local $R^2$ values.

We present here the results of the GWR models used to downscale BIO1 and BIO12, as these two variables were ultimately used for modelling the alien species distribution.

Geographically weighted regression models used to generate the topoclimatic layers scored high predictive performance for both mean temperature and annual precipitation. In particular, the GWR for mean temperature showed a strong agreement between observed and predicted mean temperatures over the whole island, with local $R^2$ values ranging from 0.974 to 0.986 (median = 0.98) (Figure S2.1). Despite a wider range of local $R^2$ values (0.226-0.982, median = 0.88), the GWR for annual precipitation also accurately predicted the precipitation pattern across most of the island. The lowest agreement between observed and predicted precipitation occurred in a restricted “bad-fit” area at the north-east side of Possession Island (Figure S2.1).

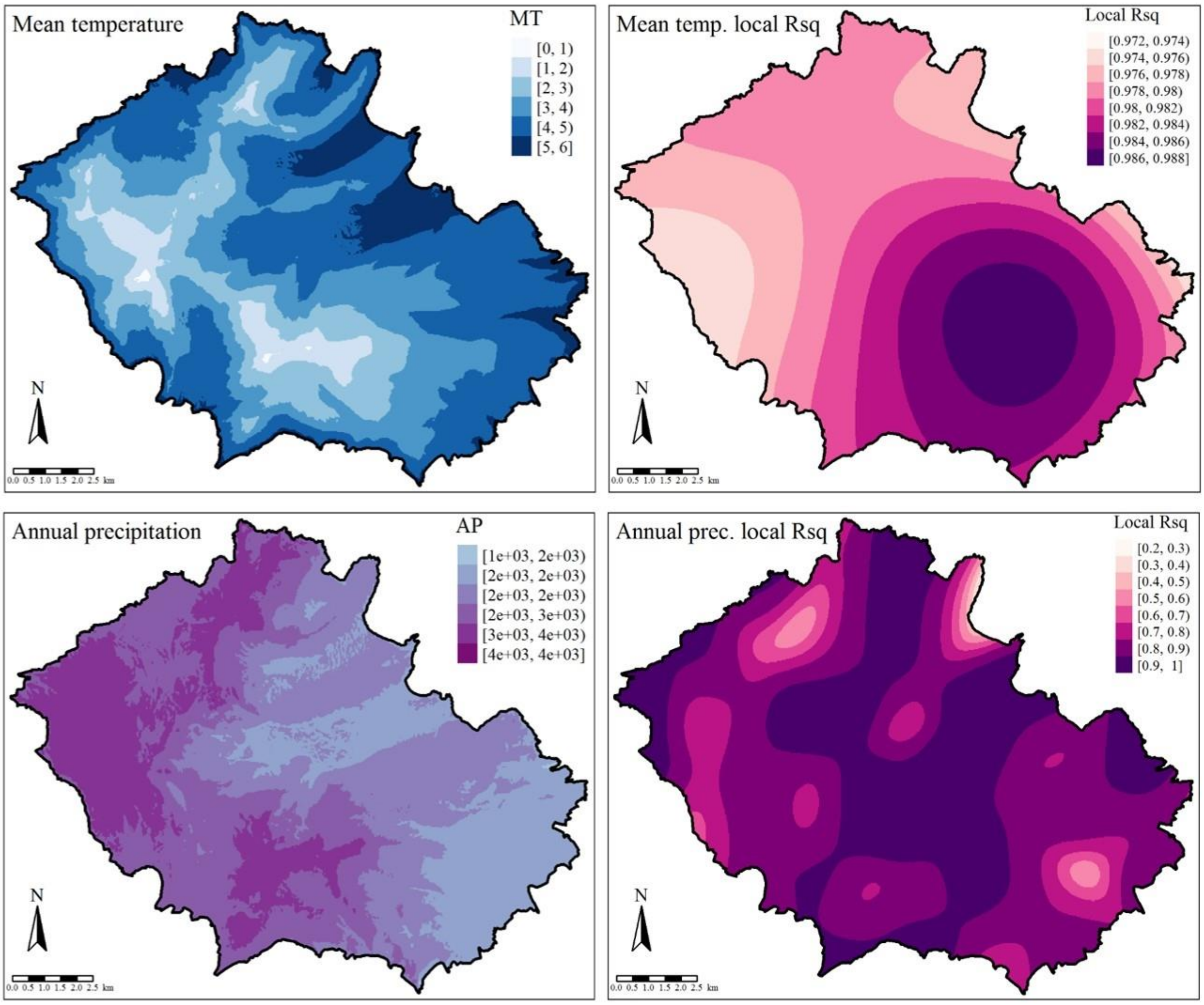


*Figure S2.1 – On the left panel, (30-m resolution) maps of the mean temperature (top) and annual precipitation (bottom) obtained through the GWR-based downscaling of the (1-km resolution) CHELSA mean temperature (BIO1) and annual precipitation (BIO12) layers. On the right panel, local $R^2$ maps showing the predictive accuracy of the GWR models. Overall, the GWR model for mean temperature (top) accurately predicted temperatures across the whole study area, while the GWR model for annual precipitation (bottom) evidenced a small "bad-fit" area at the north-east side of Possession Island. MT: mean temperature; AP: annual precipitation.*

### Appendix S3

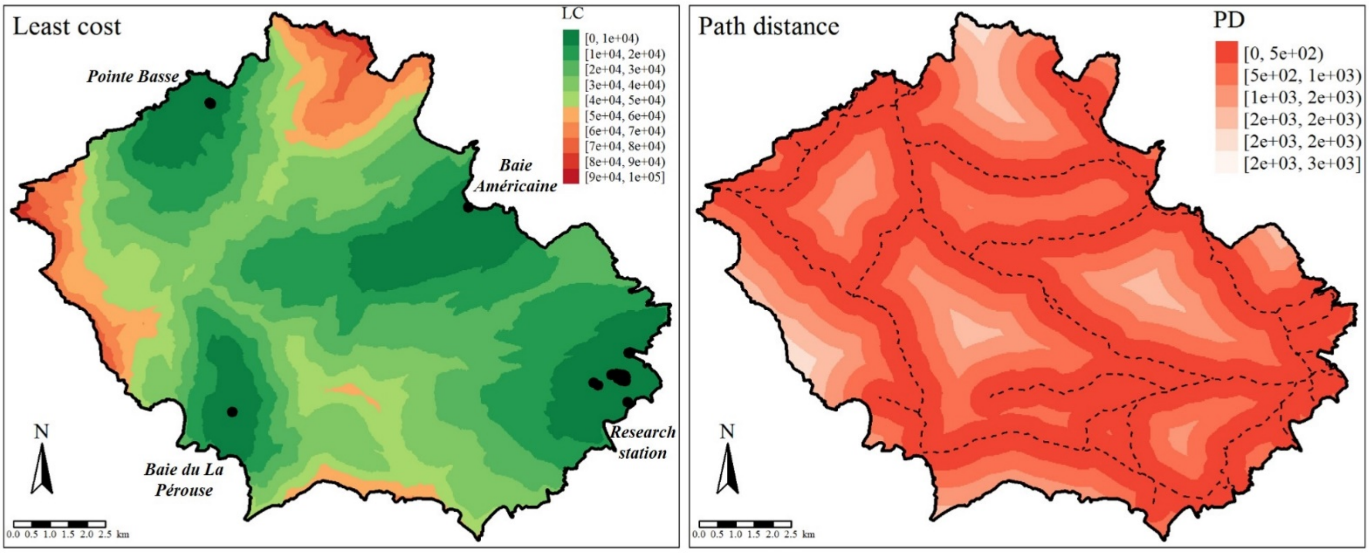


*Figure S3.2 – Left panel: map of the least cost distance from human settlements (black dots) to any location on Possession Island (LC: least cost). Right panel: map of the distance from hiking paths (dashed black lines) to any location on Possession Island (PD: path distance).*

### Appendix S4

*Environmental matching of presence/absence data*

For calibration and validation purposes, we divided the species-specific datasets in training and testing partitions reflecting all available environmental conditions on Possession Island through a systematic sampling of the species occurrence data in the environmental space. First, we summarized the environmental space by reducing the information represented by all available environmental, topographic and topoclimatic variables (see Appendix S2) through a principal component analysis (PCA) and keeping the first three components. Second, using the "eSample" function from the "iSDM" R package (Hattab et al., 2017), we identified 663 environmental pixels allowing an adequate sampling of the 3D-PCA space. We then projected the species occurrence data onto the environmental space, so as to identify a unique set of closest environmental pixels-occurrence matching pairs using the same approach described in Lenoir et al. (2010). As a last step, we selected occurrence data whose Euclidean distance from the matched environmental pixel was lower than the mean Euclidean distance of all matching pairs in the 3D-PCA space and included them in the training dataset, while we fed the testing dataset with all remaining occurrence data (Figure S4.3). The process was repeated separately for presence and absence data (for each species) to obtain a sufficiently large sample size to train the models. The final sample sizes used to train and test the single-SDMs are reported in Table S4.3.

A GIF showing an animated example of matched presences (*Cerastium fontanum*) in the environmental space is attached to the manuscript.

*Table S4.3 – Number of presence and absence observations constituting the species-specific training and testing datasets.*

| *Species* | *Training dataset (Pres. - Abs.)* | *Testing dataset (Pres. - Abs.)* |
|---|---|---|
| *Cerastium fontanum* | 421 - 409 | 690 - 439 |
| *Poa annua* | 413 - 415 | 444 - 613 |
| *Poa pratensis* | 179 - 434 | 140 - 1001 |
| *Sagina procumbens* | 380 - 428 | 392 - 664 |
| *Juncus bufonius* | 81 - 439 | 77 - 1065 |
| *Stellaria alsine* | 79 - 436 | 58 - 1037 |

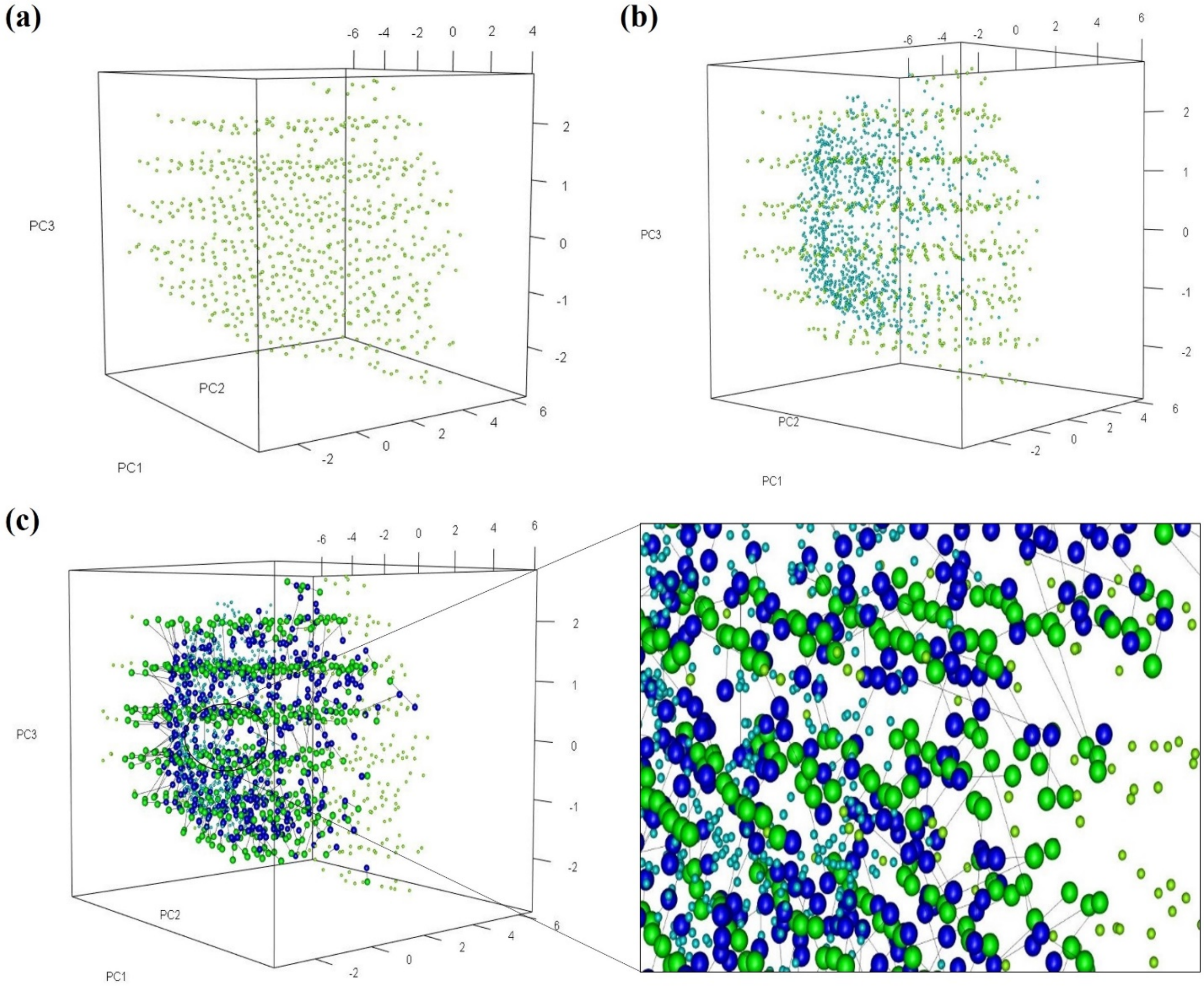


*Figure S4.3 – Graphical representation of the environmental matching procedure. Panel (a) represents the 663 environmental pixels (light green spheres) identified by the "eSample" function from the R package "iSDM" (for further details see Hattab et al., 2017). Panel (b) represents both the environmental pixels (light green spheres) and the occurrence data (light turquoise spheres, in this case representing presences of Cerastium fontanum). Panel (c) represents the best matching environmental pixels-occurrence pairs (i.e. environmental pixels-occurrence pairs lying at a distance which is below the mean Euclidean distance of all matching pairs). Solid lines connect matched environmental pixels-occurrence pairs, which appear as spheres bigger than the unmatched elements. The bottom-right picture provides an enhanced view into the environmental space.*

**Appendix S5**

Regression coefficient estimates and 95% likelihood profile-based confidence intervals for regression parameters of the low-spread (a) and high-spread (b) single-species distribution models. Likelihood profile-based confidence intervals were computed using the function "conf.int" after loading the R package "MASS" (Venables & Ripley, 2002). Superscript $^2$ identifies quadratic terms. MT: mean temperature; AP: annual precipitation; LC: least cost; PD: path distance.

**(a)**

| | *Poa pratensis* | | | *Juncus bufonius* | | | *Stellaria alsine* | | |
|---|---|---|---|---|---|---|---|---|---|
| **Predictors** | Estimate | 2.5% | 97.5% | Estimate | 2.5% | 97.5% | Estimate | 2.5% | 97.5% |
| **MT** | 0.728 | 0.092 | 1.409 | 92.587 | 20.126 | 201.991 | 0.214 | -0.528 | 0.993 |
| **MT$^2$** | / | / | / | -71.870 | -124.144 | -34.527 | / | / | / |
| **AP** | -0.005 | -0.007 | -0.004 | -0.005 | -0.007 | -0.003 | -30.406 | -47.904 | -16.631 |
| **AP$^2$** | / | / | / | / | / | / | 16.416 | 6.864 | 25.755 |
| **LC** | -0.00004 | -0.00006 | -0.00001 | -0.0001 | -0.0002 | -0.0001 | -0.000029 | -0.000058 | 0.000002 |
| **PD** | -0.002 | -0.003 | -0.001 | -0.002 | -0.004 | -0.001 | -0.0018 | -0.0032 | -0.0005 |

**(b)**

| | *Poa annua* | | | *Sagina procumbens* | | | *Cerastium fontanum* | | |
|---|---|---|---|---|---|---|---|---|---|
| **Predictors** | Estimate | 2.5% | 97.5% | Estimate | 2.5% | 97.5% | Estimate | 2.5% | 97.5% |
| **MT** | 0.825 | 0.600 | 1.057 | 24.106 | 17.141 | 31.782 | 8.346 | 3.013 | 13.787 |
| **MT$^2$** | / | / | / | -12.256 | -19.110 | -5.982 | -5.963 | -10.286 | -1.795 |
| **AP** | 0.00038 | 0.00004 | 0.00072 | -0.00034 | -0.00070 | 0.00001 | 0.00004 | -0.00031 | 0.00038 |
| **AP$^2$** | / | / | / | / | / | / | / | / | / |
| **LC** | 0.000012 | 0.000002 | 0.000022 | 0.000005 | -0.000004 | 0.000015 | 0.00002 | 0.00001 | 0.00003 |
| **PD** | -0.000408 | -0.000818 | -0.000004 | -0.0006 | -0.0011 | -0.0002 | -0.0003 | -0.0007 | 0.0001 |

### Appendix S6

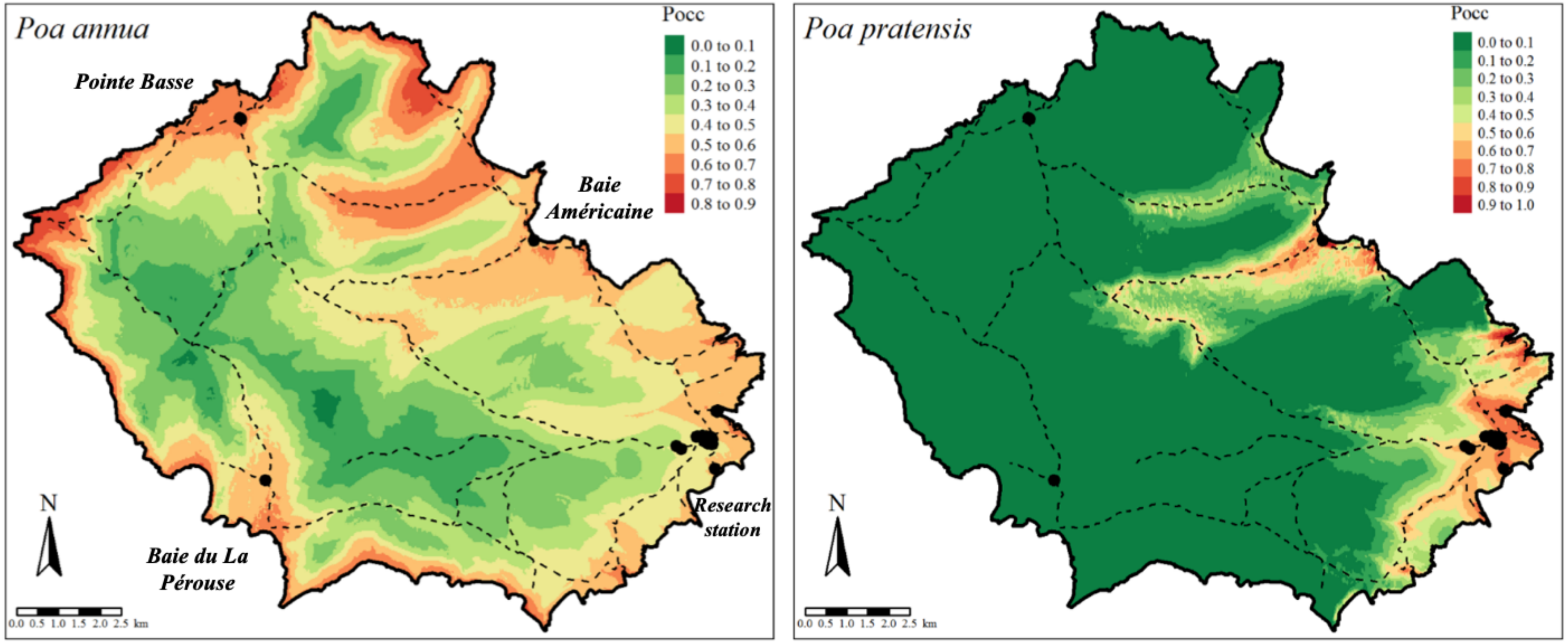


*Figure S6.4 – Predicted occurrence of Poa annua and Poa pratensis. Pocc = occurrence probability. Dashed lines represent hiking paths, while black dots represent human settlements.*

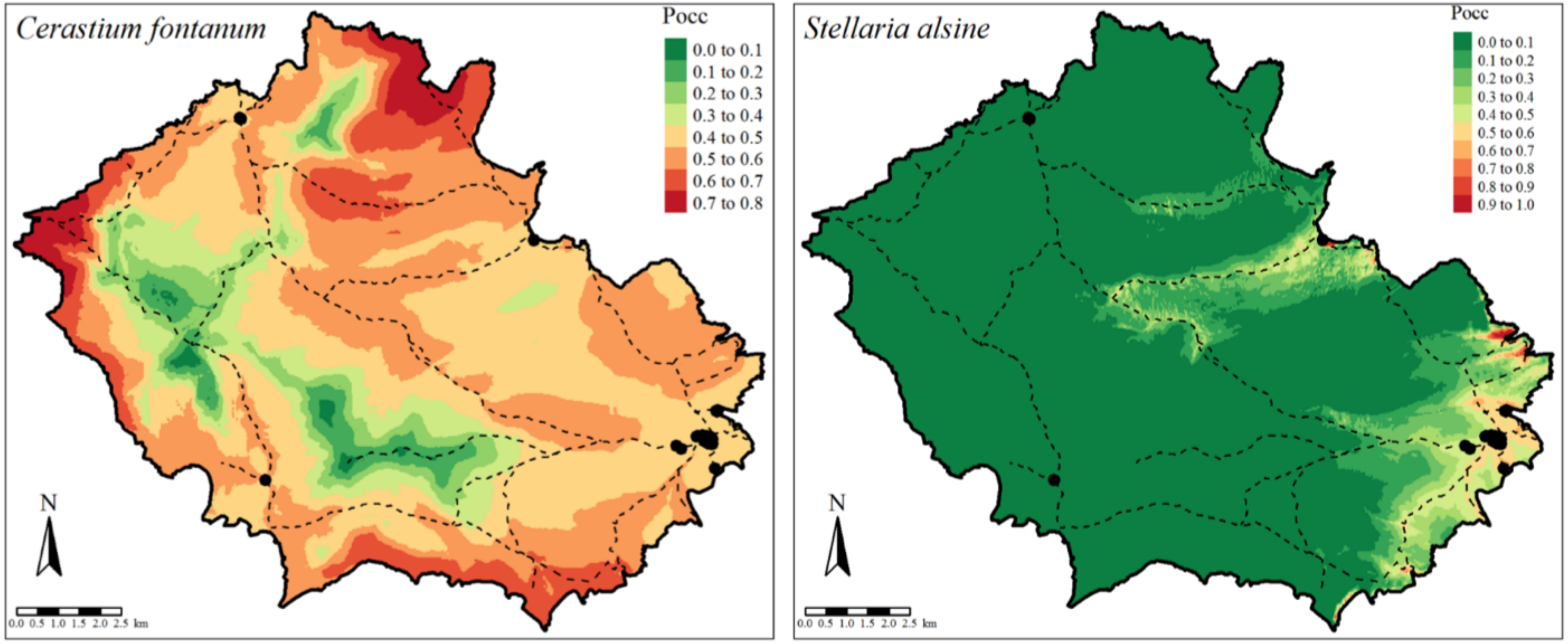


*Figure S6.5 – Predicted occurrence of Cerastium fontanum and Stellaria alsine. Pocc = occurrence probability. Dashed lines represent hiking paths, while black dots represent human settlements.*

**Appendix S7**

*Association between species features and anthropogenic propagule pressure*

To measure the relative importance of human-related variables in determining alien species occurrence in the single-SDMs, we generated a set of sub-models including all possible combinations of topoclimatic and human-related predictors (function “dredge”, R package “MuMIn”; Barton, 2019) and then summed the Akaike weights across all sub-models in which path distance and/or least cost were included (Burnham & Anderson, 2002). In particular, we summed the Akaike weights of the sub-models where these predictors had negative coefficients (i.e. where the species occurrence probability decreased further away from human facilities), as we were specifically interested in investigating the dependence on human presence. The sum of Akaike weights provides an easily interpretable measure of variable importance: it ranges from 0 to 1, with a high value for a given variable indicating its high importance relative to the others (Burnham & Anderson, 2002). We then graphically related the sum of weights for path distance and least cost to the species-specific values of the functional traits. The aim was to look for functional traits associated with low sum of weights of the human-related variables, thus highlighting plant characteristics conferring greater invasiveness.

The trends of the relationship between species functional traits and the two human-related variables (path distance and least cost) were quantitatively similar. Residence time, life history, vegetative reproduction and plant height appeared to be the plant features most related with the alien plant species dependence on human-related propagule pressure (Figure S7.6 and S7.8). In particular, old residents were associated with lower sum of weights for path distance and least cost than new residents. Similarly, annuals and species reproducing sexually had a weaker relationship with path distance and least cost than, respectively, perennials and species reproducing both sexually and vegetatively. Finally, short plants (*Sagina procumbens*, *Cerastium fontanum* and *Poa annua*) appeared to be less conditioned on human-related variables than tall ones (*Poa pratensis*, *Stellaria alsine*, *Juncus bufonius*).

On the contrary, seed- and leaf-related traits (leaf area, specific leaf area, seed dry mass and seed number per plant) did not show any association with human-related variables (Figure S7.7 and S7.9).

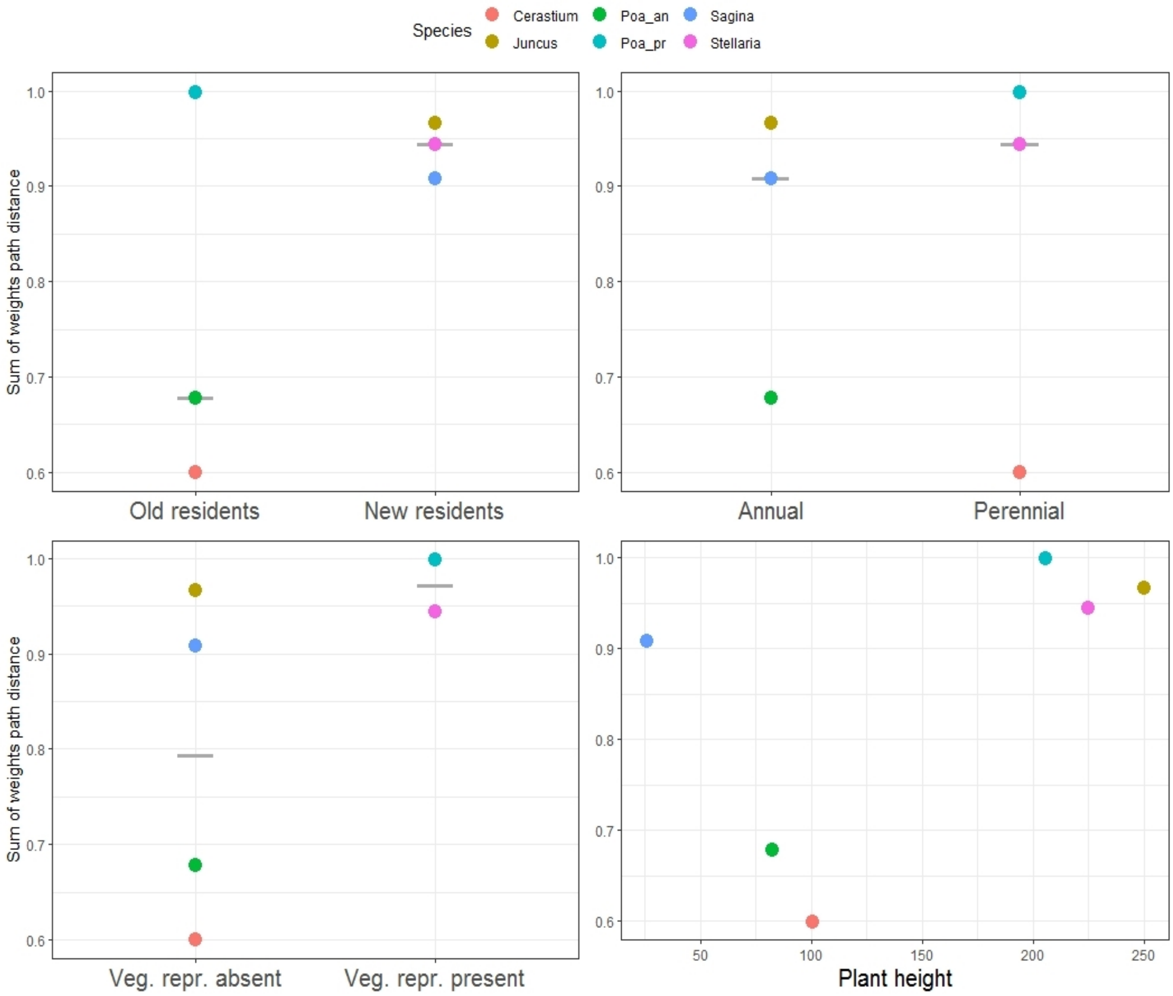


*Figure S7.6 – Relationship between sum of weights of path distance and species' residence time (top-left), life history (top-right), vegetative reproduction (bottom-left) and plant height (bottom-right). Grey crossbars represent median values.*

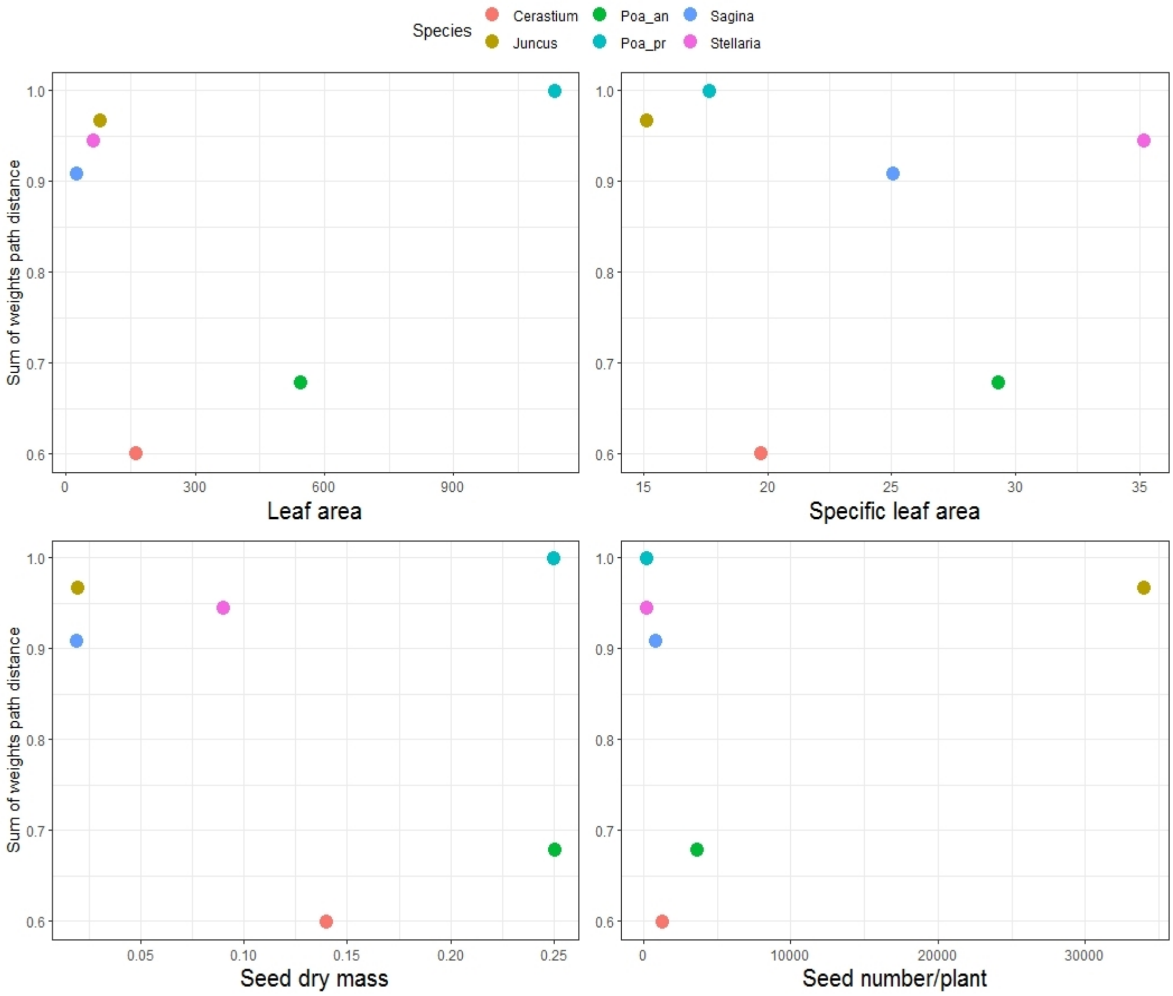


*Figure S7.7 – Relationship between sum of weights of path distance and species' leaf area, specific leaf area, seed dry mass and seed number per plant. No substantial relationship was observed between path distance and these species' traits.*

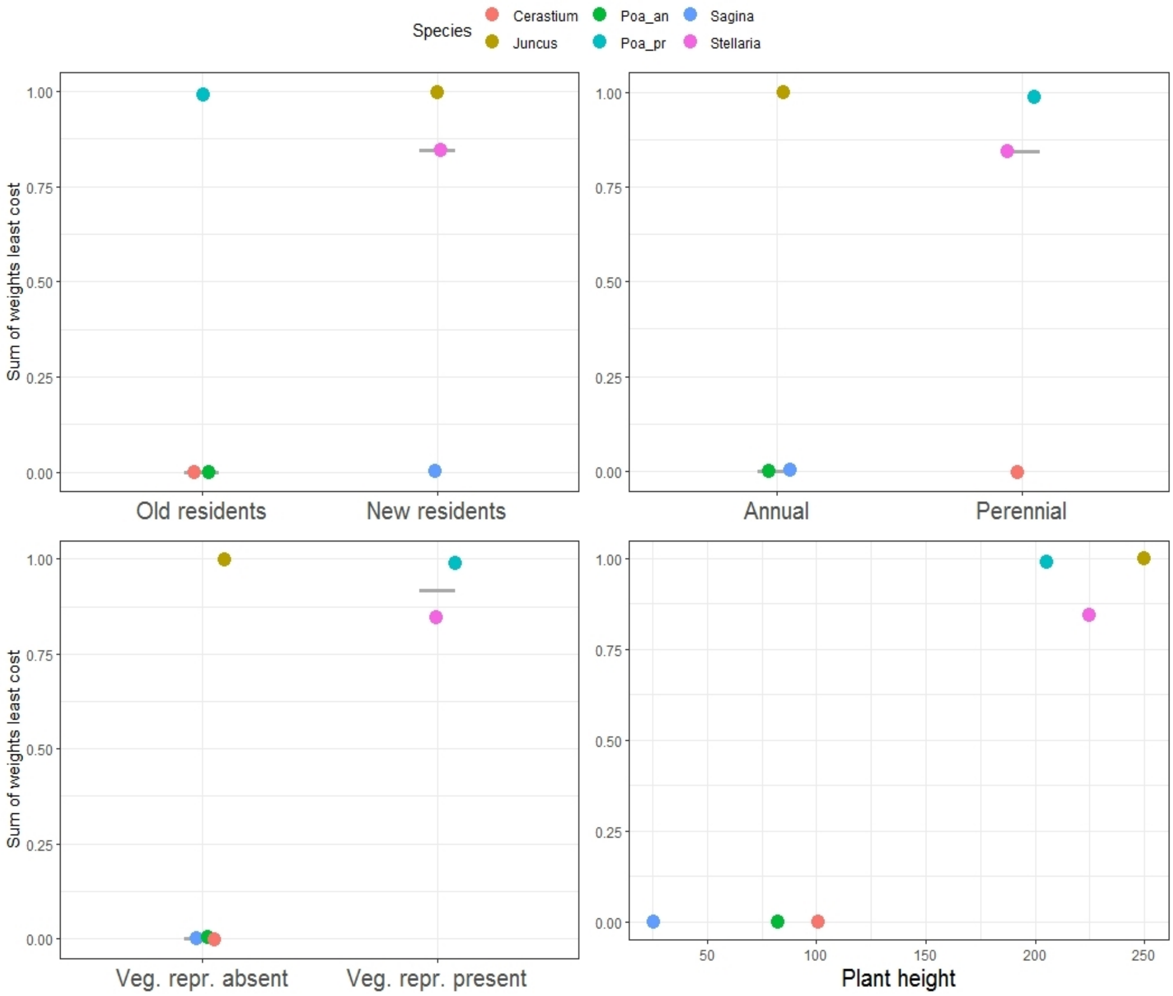


*Figure S7.8 – Relationship between sum of weights of least cost and species' residence time (top-left), life history (top-right), vegetative reproduction (bottom-left) and plant height (bottom-right). The relationships observed between least cost and these species characteristics were similar to those observed for path distance. Grey crossbars represent median values.*

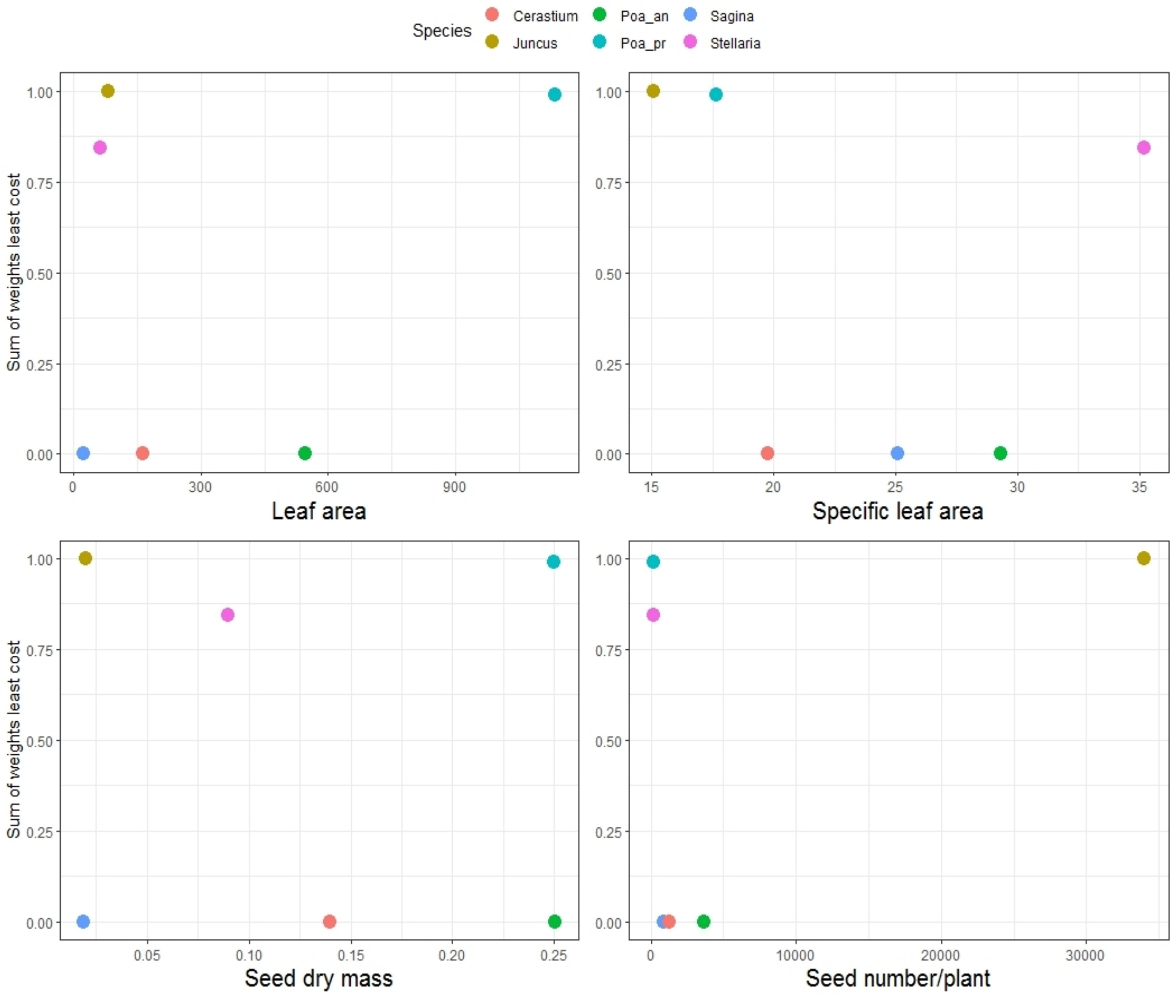


*Figure S7.9 – Relationship between sum of weights of least cost and species' leaf area, specific leaf area, seed dry mass and seed number per plant. No substantial relationship was observed between least cost and these species' traits.*

*Interaction between species features and anthropogenic propagule pressure*

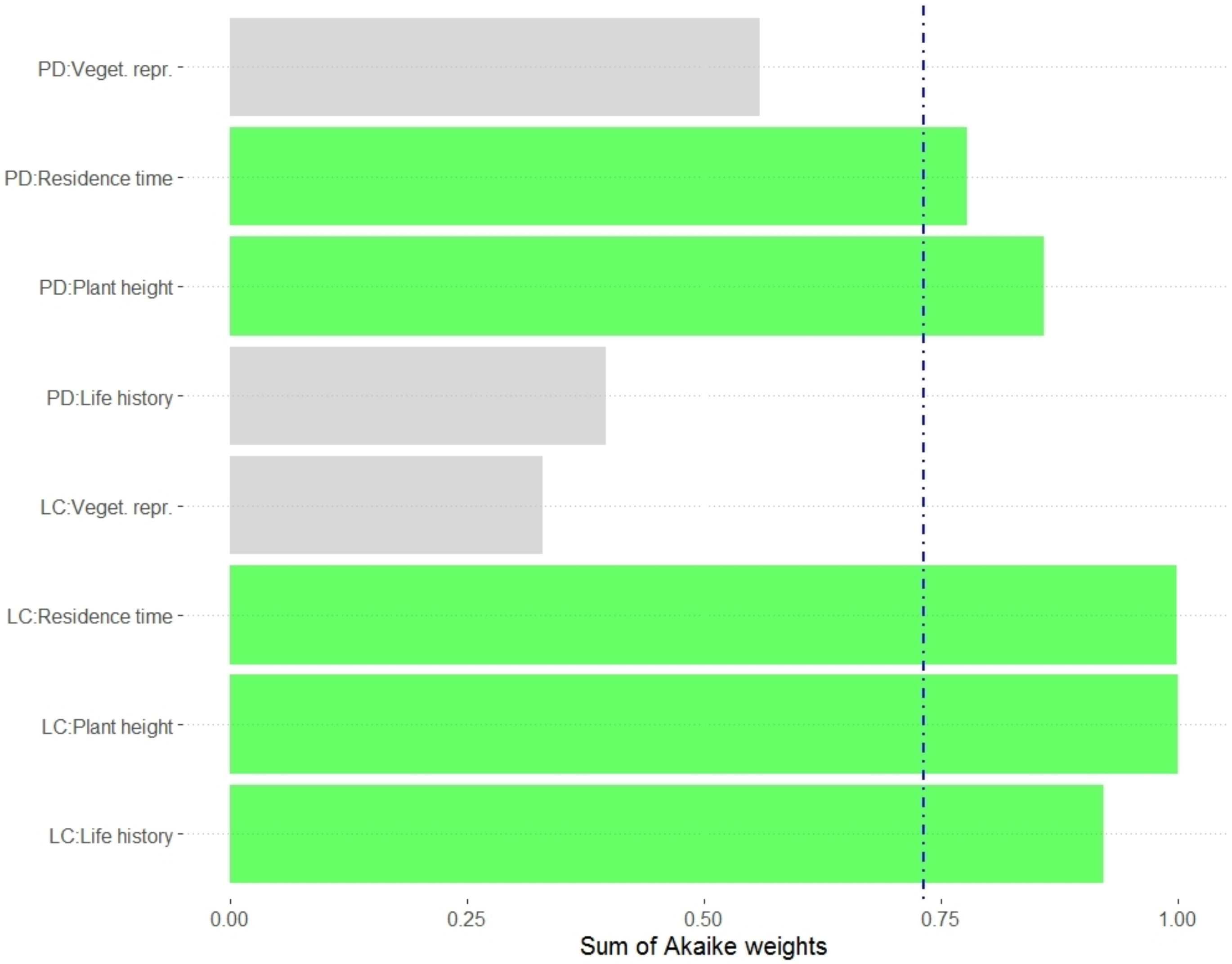


*Figure S7.10 – Sum of weights of the interaction terms between species' traits and human-related variables included in the multi-species distribution model (all other predictors were always retained in the fitted sub-models). Terms whose evidence ratio was above the threshold of 2.72 (corresponding to the sum of Akaike weights value represented by the blue dot-dash line) were retained in the most parsimonious multi-SDM and are highlighted in green, while excluded terms are highlighted in grey. PD: path distance; LC: least cost.*

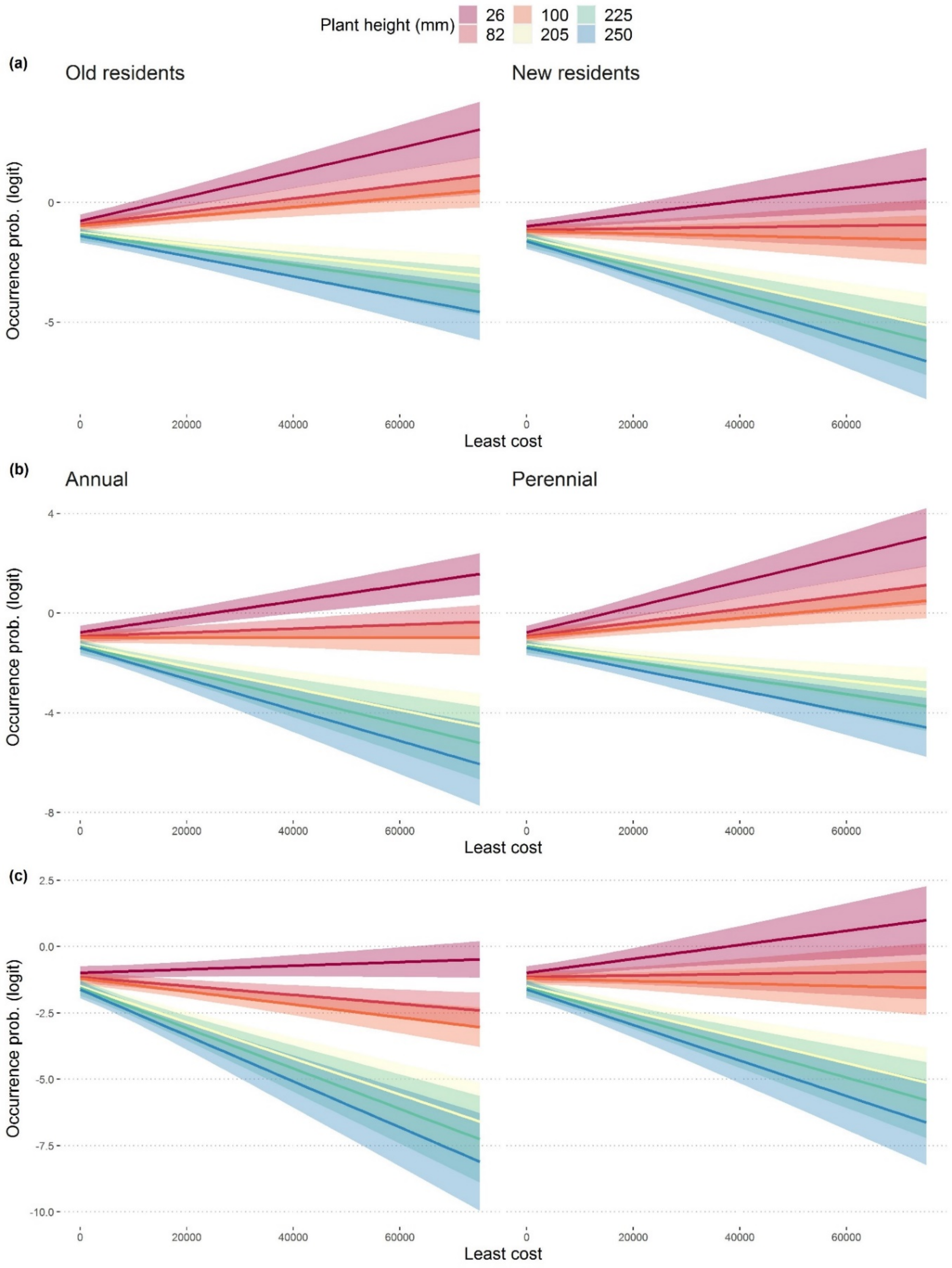


*Figure S7.11 – Effect of the interaction between least cost and plants' features (plant height, residence time and life history) on alien species occurrence probability (logit scale). Panel (a): effect of the least cost-plant height interaction on (perennial) old and new residents occurrence probability. Panel (b): effect of the least cost-plant height interaction on (old residents) annual and perennial alien species occurrence probability. Panel (c): effect of the least cost-plant height interaction on (new residents) annual and perennial alien species occurrence probability. All plots are reported at the logit scale.*
