## Supplementary figures and images for "Once upon a time in the far south: Influence of local drivers and functional traits on plant invasion in the harsh sub-Antarctic islands"

### Environmental matching

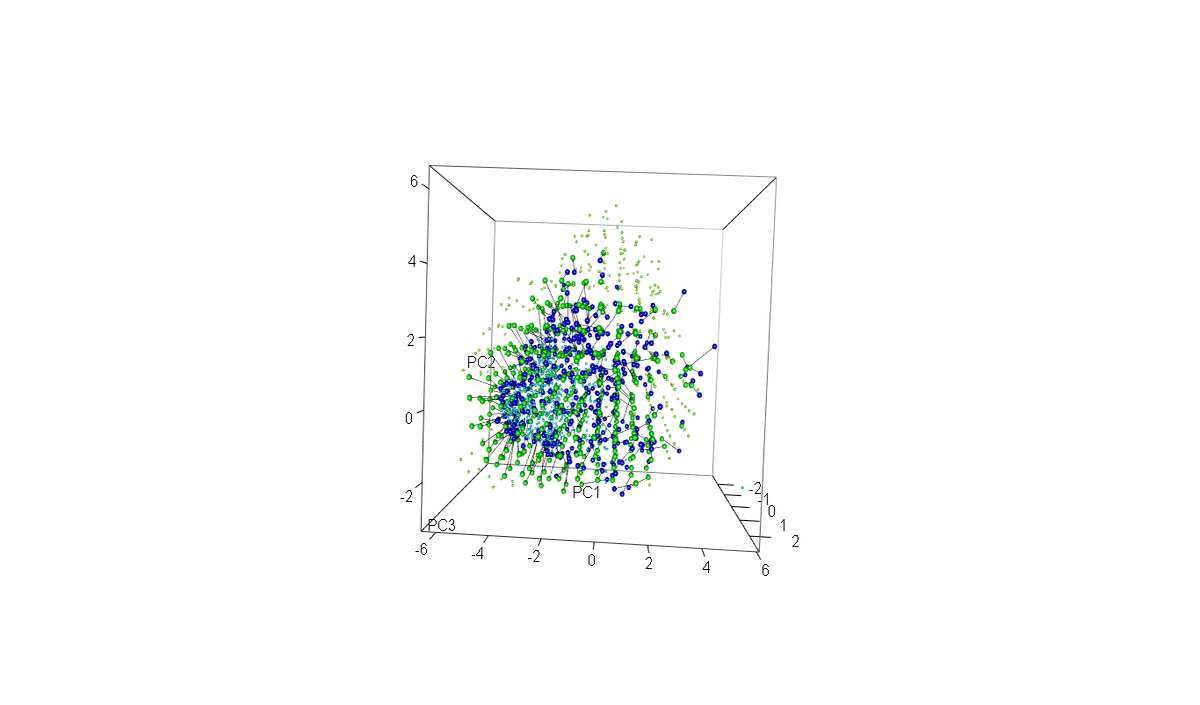
